# In situ profiling of nanoscale displacements uncovers mechano-architectural predictors of osteoarthritis emergence

**DOI:** 10.64898/2025.12.04.692296

**Authors:** Aikta Sharma, Lucinda AE Evans, Lucie E Bourne, Jishizhan Chen, Alissa L Parmenter, Joseph Brunet, Kamel Madi, Sebastian Marussi, Paul Tafforeau, Andrew A Pitsillides, Peter D Lee, Katherine A Staines

## Abstract

Mechanical and anatomical interplay between the distinct tissues of the knee joint is essential for maintaining functional integrity during healthy ageing and contributes to the mechanisms that drive osteoarthritis (OA). In this study, we investigate how age- and disease-associated alterations in joint anatomy influence load transmission and tissue-level strain distribution. Using full-field synchrotron X-ray computed tomography coupled with digital volume correlation, we hierarchically characterised in situ nanoscale strains generated in response to mechanical loading across the tibial epiphysis. Our findings show that greater compressive strains accumulate in the articular condyle of male OA-prone (STR/Ort) epiphyses. Finite element modelling further demonstrated that these strain concentrations are associated with reduced load-bearing capacity, which arise from architectural differences localised to the subchondral bone plate. By coupling high-resolution imaging with computational modelling, our work provides new insights into how structural-function changes to joint anatomy contribute to the initiation and progression of mechanically driven OA. Our approach offers a means to identify early imaging biomarkers prior to OA diagnosis and has potential for monitoring interventions aimed at preserving joint mechanics while promoting healthy joint ageing.

## Introduction

The tibial epiphysis is a key structural and functional component of the knee joint that is central to the maintenance of joint integrity and efficient load transfer. While the tissues of the tibial epiphysis, namely the articular cartilage, subchondral plate (SCP) and trabecular bone are physiologically and mechanically distinct (1), how these act in concert to maintain joint health throughout life remains poorly understood. Osteoarthritis (OA) is the most prevalent age-related degenerative joint disease, affecting approximately 8% of the global population (2, 3). Despite its widespread burden, there are currently no disease-modifying therapeutics or predictive biomarkers for OA with end-stage disease often necessitating costly and invasive joint replacement procedures (4, 5). A major barrier to therapeutic progress lies in the incomplete understanding of how mechanical and structural interplay within the tibial epiphysis contributes to the initiation and progression of OA.

In health, the articular cartilage relies on a dynamic equilibrium of factors involved in its homeostasis to ensure life-long integrity that is critical for healthy joint function (6). The underlying SCP functions as a key biomechanical interface to absorb, redistribute and transmit loads arising within the joint while influencing the mechanobiological environment of the overlying cartilage and underlying trabecular bone (7–9). In OA, however, the progressive loss of articular cartilage is accompanied by aberrant thickening of the SCP, perturbing this mechanical dependence and perpetuating cycles of abnormal bone remodelling and ineffective protection of the articular cartilage (10, 11). Indeed, a major challenge in understanding the biological, physical and mechanical crosstalk between constituent tissues of the epiphysis is critical to understanding OA pathophysiology prior to marked cartilage loss. To unravel these structure–function relationships, the development and application of enabling imaging approaches capable of resolving the microstructure of joint tissues while simultaneously characterising their mechanical behaviours is crucial.

Our previous work established synchrotron X-ray computed tomography (sCT) as a powerful modality for visualising intact murine joints across hierarchical length scales under habitual weight-bearing load, offering superior resolution, contrast, and imaging speed compared with laboratory-based micro-computed tomography (microCT) (12). sCT has since been successfully applied to characterise joint tissue microstructure, but not mechanical behaviour, in several animal models (13–16) and in human hip, knee, foot, and ankle joints (14, 17–21). Meanwhile, digital volume correlation (DVC) has emerged as a transformative computational method for tracking voxel-level greyscale patterns in sequential volumetric images acquired under load (22, 23) with integration with CT modalities now permitting the quantification of full-field three-dimensional strain in skeletal and osteochondral tissues (24–30). When combined with finite element (FE) modelling that has been calibrated using DVC-derived strains, it is possible to estimate local mechanical properties therefore enabling conclusive evaluation of load bearing behaviours (25, 31–33).

Herein, we leveraged the Extremely Brilliant Source (EBS) upgrade at the European Synchrotron Radiation Facility to perform high-resolution phase-contrast sCT imaging of intact, in situ loaded fresh murine knee joints from mice prone to developing spontaneous OA (STR/Ort) and parental, healthy ageing controls (CBA, Fig. 1A; (34–36)). By coupling sCT with DVC analysis, we examined how physiologically relevant compressive loads are accommodated across the entire tibial epiphysis during normal ageing observed in CBA mice, and how these patterns are disturbed in the knee joints of STR/Ort mice predisposed to load-induced failure and OA, akin to that in humans (36). By integrating FE simulation and microarchitectural anatomical characterisation with our approach, we discovered how variation in the SCP and trabecular bone within the tibial epiphysis influences local strains generated during healthy ageing, and how the loss of this mechanical integrity triggers the emergence of joint degeneration in OA.

**Figure 1.**
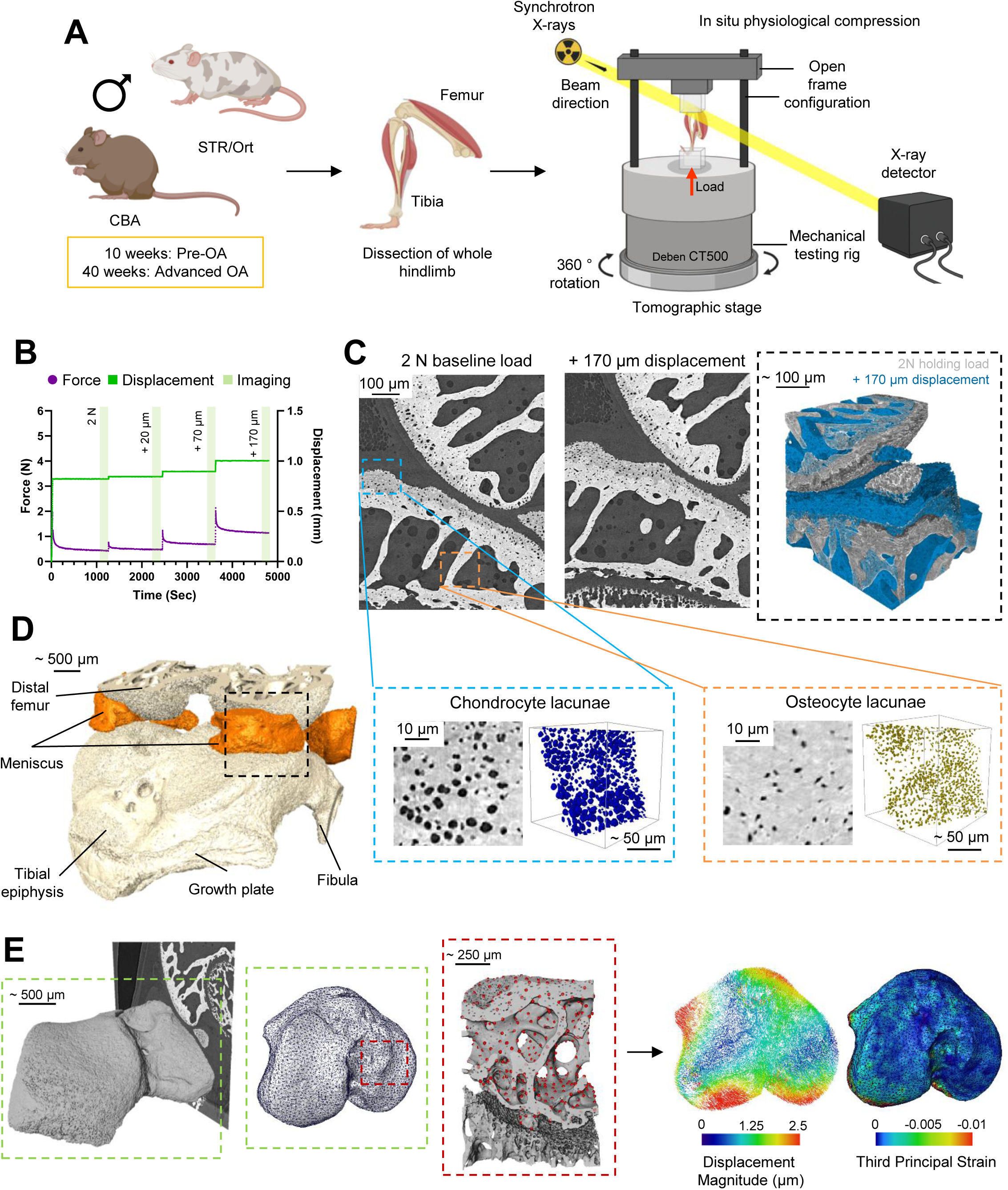
Characterisation of load-induced compressive strain in the epiphyses of healthy CBA and OA-prone STR/Ort knee joints with phase-contrast sCT and DVC. Schematic of methodology showing isolation of hindlimbs from 10- and 40-week-old male CBA and STR/Ort mice and mounting in a Deben CT500 in situ compression stage with a bespoke open frame, for in situ mechanical compression testing and sCT (A). Representative force (purple line) and displacement (green line) readings with acquisition of sCT (green rectangle) images performed after the application of a 2 N baseline load and three displacement-controlled compression steps of 20 µm, 70 µm and 170 µm (B). Sagittal view of sCT images of the medial tibial epiphysis following the application of the 2 N baseline load (C, left) and in response to 170 µm compressive displacement (C, middle) are shown in 3D (C, right; grey, 2 N baseline load; blue, 170 µm displacement). Zoomed sCT images and 3D renders of epiphyseal microstructure are shown for chondrocyte lacunae (blue box) and osteocyte lacunae (orange box) within epiphyseal SCP. 3D rendering of the CBA joint with anatomical regions labelled (D). 3D rendering of the tibial epiphysis, including growth plate bridges, segmented by a region-growing algorithm and closure of chondrocyte and osteocyte lacunae (E, left) were used for the generation of finite element tetrahedral meshes (E, green hatched box) with corresponding point cloud nodes overlaid on epiphyseal microstructure (E, red hatched box) shown and exploited used for the calculation of displacement and strain magnitude by DVC (E, right). Schematic shown in A was created using biorender.com.

## Results

### Elevated load-induced strains arise prior to OA emergence in the STR/Ort tibial epiphysis

To resolve in situ epiphyseal biomechanical strains by DVC, we first upgraded a regime for recapitulating knee joint loading in intact murine hindlimbs during typical locomotion compatible with imaging by phase-contrast enhanced sCT (37) (Fig. 1A). In line with previous studies, an initial 2 N load was applied to fix hindlimbs in a flexed position (38) (Fig. 1B). As anticipated, joints exhibited viscoelastic stress relaxation under this 2 N ‘baseline’ uniaxial load and were thus sCT imaged after a period of 15 minutes, when force stabilisation was attained. Successive sCT images were captured at three displacement-controlled load steps of 20, 70 and 170 µm, which produced peak forces of 1.0-3.5 N and relaxed/imaged forces of 0.5-1.5 N (52-63% lower) across age groups and genotypes (Supplementary Fig. 1A and B).

Inspection of sCT images at baseline load confirmed expected age and genotype phenotypes; 10- and 40-week-old CBA showed normal joint microarchitecture, while STR/Ort mice displayed the expected progressive osteoarthritic changes to joint microarchitecture, hallmarked by subchondral bone sclerosis, osteophytosis and joint space narrowing at 40 weeks of age (Supplementary Fig. S2A). Ensuing histological evaluation (Supplementary Fig. S2B and C), through OARSI scoring, further confirmed conservation of tibial articular cartilage integrity in 10-week-old STR/Orts and all CBA mice, as well as marked lesions on medial (p=0.01) and lateral (p=0.005) condyles in STR/Ort at 40 weeks of age, indicative of OA progression (Supplementary Fig. S2D).

The enhanced resolution of sCT images affords FE tetrahedral mesh creation from segmentations of reconstructed epiphyseal volumes (Fig. 1C-E). Nodal displacements in each mesh were estimated by correlating greyscale intensity variations (baseline vs. loaded) and the resultant displacement fields used for full-field strain computation by the global DVC approach. This yielded measurement accuracy for displacement of 18 nm in the loading direction (z-axis) and a precision of 78 nm, culminating in mean strain accuracy of 53 µE and 221 µE mean strain precision.

Herein, the subsequent quantification of displacement-induced compression (E3, third principal) and tension (E1, first principal) in CBA and STR/Ort epiphyses enabled exploration into whether physiological load induces divergent deformation patterns in healthy and OA-prone joints. This exposed a near-uniform low compression across the SCP of 10-week-old CBA epiphyses (Fig. 2A, left, top), with raised strains transmitted to growth plate regions (Fig. 2A, left, bottom). In contrast, age-matched STR/Ort showed subtle elevations in compression (compared to CBA), with strain accumulation evident in the lateral SCP and growth plate regions (arrowheads). Examination of older 40-week-old CBA showed that compression levels were homogenous and effectively equivalent to those seen across the SCP in young CBA epiphyses (Fig. 2A, right). In stark contrast, loading generated high compressive strain foci at regions of the SCP in 40-week-old STR/Ort epiphyses, where growth plate surfaces accumulated greater compression levels, indicative of a markedly more deformable epiphysis (Fig. 2A, right, bottom).

**Figure 2.**
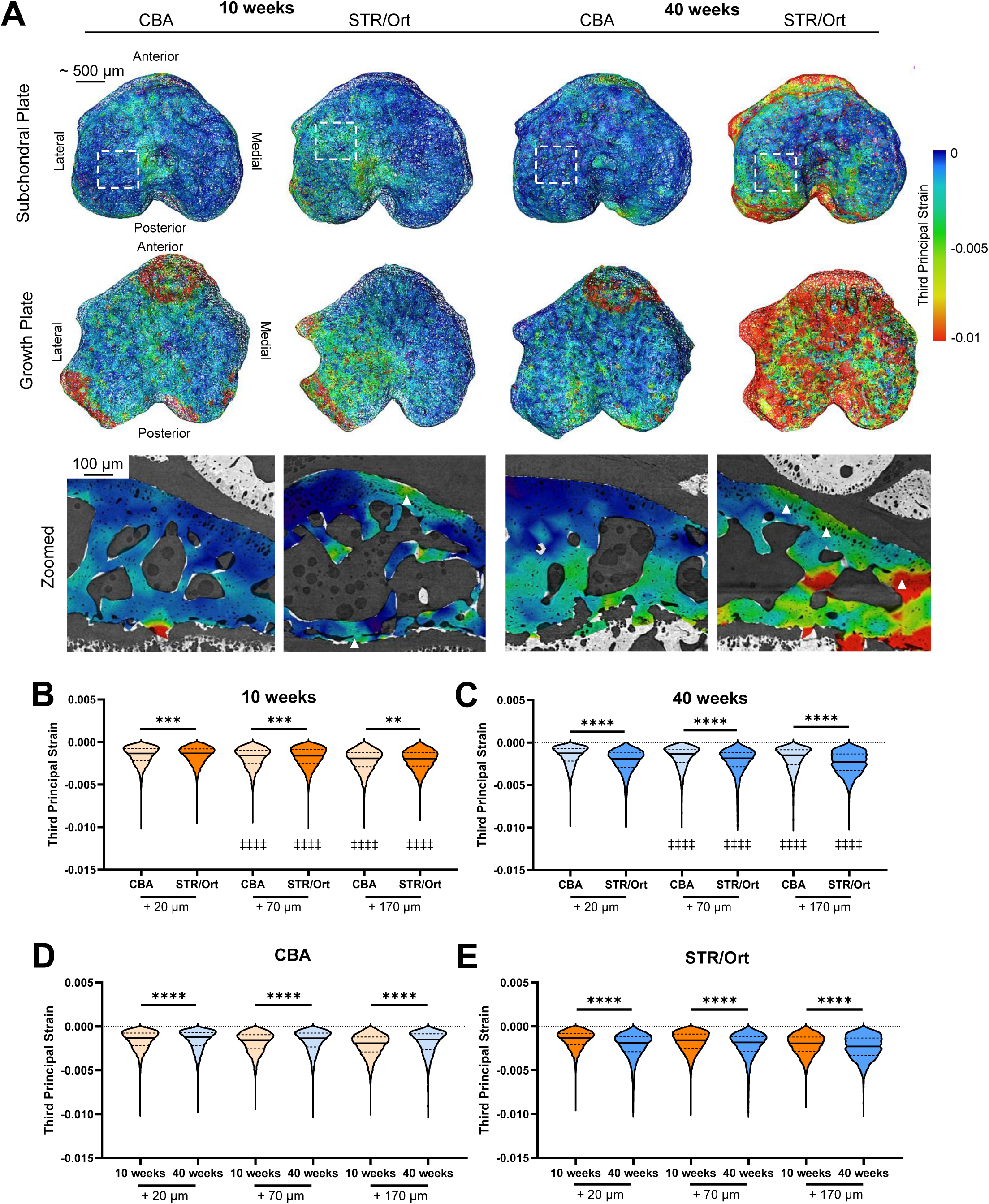
Emergence of OA pathology in STR/Ort mice is associated with the regionalised accumulation of compressive strains. DVC-computed strains are superimposed on finite element tetrahedral meshes of 10-and 40-week-old (right) CBA and STR/Ort tibial epiphyses, viewed from the SCP (A, top row) and from the growth plate (A, bottom row) show localisation of high (red) and low (blue) compressive strains. In CBAs, high magnitude compressive strains are confined to growth plate regions of the tibial epiphyses in both 10-and 40-week-old animals. In STR/Orts, compressive strains of relatively higher magnitude are evident across the SCP and growth plate regions in 10-week-old animals which further increase in magnitude in 40-weeks-olds in line with OA progression. Zoomed view of compressive strains overlaid on sagittal sCT images (A, bottom) highlight high strain accumulation within distal epiphysis in CBAs and in the SCP and growth plate regions in STR/Orts. Quantification of the distribution of compressive strains from point cloud nodes, in response to displacement-induced load in 10-week-old animals (B) and 40-week-old animals (C) are presented in violin plots with median value indicated by the solid line, and upper and lower dashed lines representing 25^th^ and 75^th^ quartiles, respectively. Ageing is also associated with narrower compressive strain distribution in CBAs (D) while the opposite is evident in STR/Orts (E) following incremental displacement. Individual violins represent DVC-derived strains from N=4 animals/age. Statistical significance between CBAs and STR/Orts was assessed using the Kolmogorov-Smirnov test (**p<0.01, ***p<0.001 and ****p<0.0001). Statistical significance between displacement-induced load steps in CBAs and STR/Orts relative to the strains induced in response to 20 µm displacement was assessed using One-way ANOVA with Dunnett’s post-hoc test (‡‡‡‡p<0.0001).

Confirming our observations, the 20 µm displacement was found to induce significantly broader compressive strain distributions (D, cumulative distribution function) in 10-week-old STR/Ort than CBA mice (D=0.025, p=0.0004; Fig. 2B) which were equivalent and retained at 70 and 170 µm displacement (D=0.026, p=0.0002 and D=0.023, p=0.002, respectively). This divergence in compressive strain distributions in STR/Ort (versus CBA) was ∼10-fold greater at 40-weeks at all displacements (D=0.18-0.24, all p<0.0001, Fig. 2C). Across both ages, median compressive strain was, however, consistently elevated and linked to larger interquartile ranges (IQRs) in STR/Ort at successive displacements (Fig. 2B and C & Supplementary Table S1), while CBA showed only modest age-related changes (D=0.04-0.15, p<0.0001, Fig. 2D). Here, increasing displacement engendered incremental increases in median compressive strain magnitudes; these were higher in 10-week-old than 40-week-old CBA. In STR/Orts however, ageing resulted in small yet significant changes to compressive strain distribution (D=0.1-0.21, p<0.0001, Fig. 2D) and were accompanied by only slight age-related differences in median compressive strains and IQRs (Supplementary Table S1).

These findings were complimented by similar trends in tensile strain accumulation in the epiphyses of STR/Ort mice at both 10 and 40 weeks of age (Supplementary Fig. S3A-C, Supplementary Table S2). Ageing in CBA mice was linked to large changes in tensile strain (D=0.32-0.42, p<0.0001; Supplementary Fig. S3D), with median tensile strain magnitudes higher in 10-week-old CBAs than 40-week-olds (Supplementary Table S2). STR/Ort ageing was however, characterised by more marked load-induced changes to tensile strain distribution (D=0.04-0.26, Supplementary Fig. S3E), accompanied by only subtle age-related differences in median compressive and tensile strains and IQRs (Supplementary Table S1 and S2). Across both genotypes and ages, increasing displacement translated in greater compressive and tensile strain magnitudes relative to the baseline 2 N load (all p<0.0001) (Fig 2B & C and Supplementary Fig. S3B & C, respectively).Collectively, these data indicate that the greater compressive deformability of the STR/Ort tibial epiphysis precedes OA onset and may underlie increased vulnerability to load-induced failure, whereas higher tensile strains generated in CBA epiphyses upon loading appear characteristic of healthy ageing.

### Joint loading creates laterally biased deformation in the STR/Ort subchondral plate

To interrogate whether strain accumulation in STR/Ort joints is regionally confined to a particular epiphyseal compartment, DVC datasets were subdivided into distinct lateral-medial anatomical ROIs (Fig. 3A). This revealed symmetrical and uniform condylar distribution of low magnitude compressive and tensile strains in SCP and trabecular regions in 10-week-old CBA epiphyses (Fig. 3B and Supplementary Fig. 4A) that were mostly retained in 40-week-old CBA. In contrast, compressive and tensile strain distributions were laterally biased in the SCP and extracondylar cortical bone regions in 10-week-old STR/Ort, with these asymmetries becoming more marked with age in 40-week-old animals (Fig. 3B and Supplementary 4A).

**Figure 3.**
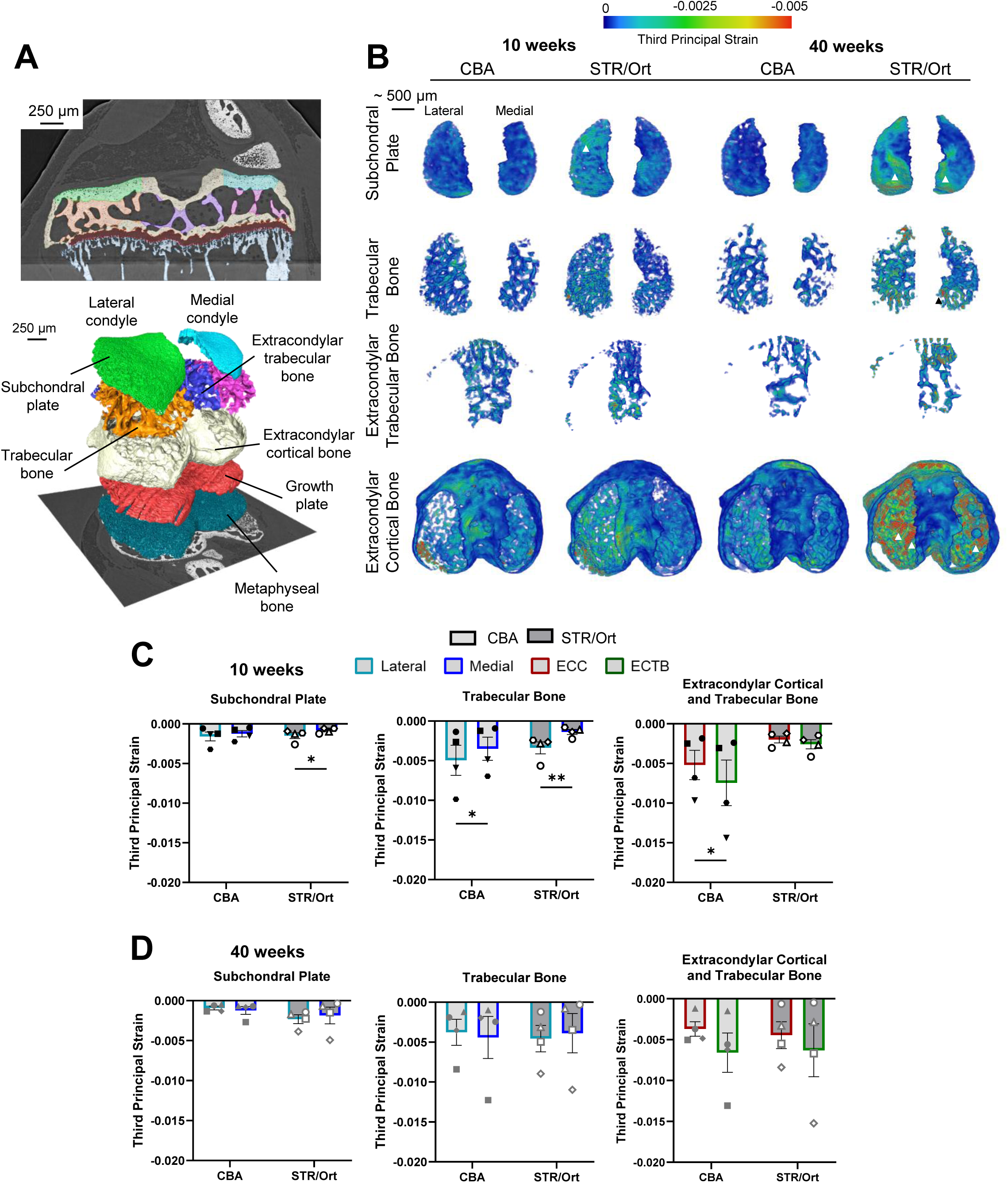
Regionalised strain quantification reveals the imbalanced distribution of compressive strain across the STR/Ort epiphyseal condyles. Segmentation of anatomical regions of interest from the lateral and medial condyles of the tibial epiphysis, including the SCP, trabecular bone, ECTB and ECC (A) shown in 2D coronal images (top) and in 3D renders (bottom). Region-specific DVC-derived compressive strain are shown for the SCP, trabecular bone, extracondylar trabecular bone and extracondylar cortical bone from 10- and 40-week-old CBA and STR/Ort mice (B). Quantification of average compressive strain in the lateral (light blue outlined bar) and medial (dark blue outlined bar) condylar compartments of the tibial epiphyses of 10-week-old (C) and 40-week-old (D) CBA and STR/Ort mice show laterally-dominant load induced condylar strains arise in STR/Orts in 10 week old animals, while latero-medial balancing of strain is evident in CBAs of both ages. Data are presented as mean ± SEM with symbols representing individual animals (N=4/age group/genotype). Statistical significance between condyles was assessed using linear mixed model analyses with Fisher’s LSD post-hoc test (C and D; *p<0.05, **p<0.01).

Confirming the spatial patterns observed in the regionalised DVC maps, quantification of average nodal strain revealed balanced compressive (Fig. 3C) and tensile (Supplementary Fig. S4B) strain magnitudes between the lateral and medial SCP in joints of 10-week-old CBA mice, while an imbalanced accumulation of compressive (p=0.03) and tensile (p=0.02) strains was evident in the lateral SCP of 10-week old STR/Ort mice. Further asymmetry was noted within the trabecular bone compartment, where strains were higher in the lateral trabecular bone compartment in both CBA (compression, p=0.03 and tension, p=0.04) and STR/Ort epiphyses (compression, p=0.007 and tension, p=0.008). In the extracondylar compartment, compressive strains were higher in the trabecular bone compared to the cortical bone in CBA epiphyses (p=0.02) whereas tensile strains were higher in trabecular than cortical bone in STR/Ort (p=0.004). Remarkably, both compressive and tensile strains were latero-medially balanced across all epiphyseal compartments in CBAs and STR/Orts at 40 weeks of age (Fig. 3D and Supplementary Fig. S4C), with no age-related effects noted. These data align with the bulk DVC, highlighting that complementary genotype and age-dependent alterations in both compressive and tensile strain patterns are already evident in the joints of 10-week-old animals.

Epiphyseal ROI subdivision into distinct anatomical compartments also allowed the material properties of each to be predicted by FE modelling; using specific meshes generated by binary epiphyseal segmentations (Supplementary Fig. S5A). Bland-Altman analysis confirmed >95% agreement of FE predictions with measured strains by DVC (Supplementary Fig. S5B and C), and strain distribution in these models showed high concordance (1-2% deviation, Supplementary Fig. S5D and E, respectively) at nodes experiencing high strain magnitudes. Regionalised DVC aligned with inverse optimisation of material properties to reveal compartment-specific variation in predicted stiffness in response to deformation across ROIs (Table 1). Application of this approach to 10-week-old CBA epiphyses showed that lateral and medial SCP and trabecular bone regions exhibited comparable moduli, and that trabecular bone was significantly less stiff than cortical bone in the extracondylar compartment (p=0.006); 10 week-old STR/Ort showed a similar pattern, with lower moduli in extracondylar trabecular bone (p=0.01) and equivalent lateral-medial behaviours. Notably, SCP stiffness was greater in 10-week-old CBAs than STR/Orts (lateral, p=0.0008 and medial, p=0.0001), whereas extracondylar trabecular bone was less stiff (p=0.03). This aligns to raised load-induced epiphyseal strains but not to the laterally biased accumulation of strain seen in STR/Ort mice.

**Table 1.** Finite element predictions of Young’s modulus and Poisson’s ratio for 10- and 40-week-old CBA and STR/Ort mice across anatomical compartments of the tibial epiphysis. Data presented as mean ± SEM for N=3 mice/age/genotype. Statistical significance was assessed using linear mixed model analysis with Fisher’s LSD post-hoc test. † denotes statistical significance between 10- and 40-week-old animals, * denotes statistical significance between CBA and STR/Ort and § denotes statistical significance between anatomical region/condyle with single, double and triple symbols corresponding to p<0.05, p<0.01 and p<0.0001, respectively.

|  |  |  | Lateral Subchondral Plate | Medial Subchondral Plate | Lateral Trabecular Bone | Medial Trabecular Bone | Extracondylar Cortical Bone | Extracondylar Trabecular Bone |
| --- | --- | --- | --- | --- | --- | --- | --- | --- |
| Predicted Modulus (GPa) | 10 weeks | CBA | 3.32 ± 0.29 | 3.8 ± 0.11 | 7.84 ± 0.55 | 8.78 ± 0.47 | 9.47 ± 0.32 | 5.38 ± 0.73<br>§§ |
| | | STR/Ort | 1.10 ± 0.39<br>*** | 0.86 ± 0.34<br>*** | 7.43 ± 1.15 | 7.86 ± 1.92 | 10.77 ± 0.69 | 7.51 ± 0.34<br>§, *** |
| | 40 weeks | CBA | 3.51 ± 0.46 | 3.96 ± 0.17 | 7.49 ± 0.33 | 5.78 ± 1.16<br>† | 10.58 ± 0.8 | 5.36 ± 0.73<br>§§ |
| | | STR/Ort | 4.13 ± 0.39<br>††† | 3.48 ± 0.15<br>††† | 1.24 ± 0.51<br>††, *** | 2.18 ± 0.57<br>††, ** | 7.83 ± 0.53<br>† | 2.89 ± 1.21<br>††, §§ |
| Predicted Poisson's Ratio (A.U.) | 10 weeks | CBA | 0.24 ± 0.01 | 0.22 ± 0.01 | 0.26 ± 0.02 | 0.27 ± 0.03 | 0.22 ± 0.01 | 0.29 ± 0.02<br>§§ |
| | | STR/Ort | 0.2 ± 0.01<br>* | 0.25 ± 0.02 | 0.24 ± 0.01 | 0.28 ± 0.02 | 0.21 ± 0.003 | 0.29 ± 0.01<br>§§ |
| | 40 weeks | CBA | 0.23 ± 0.01 | 0.21 ± 0.01 | 0.29 ± 0.02 | 0.29 ± 0.02 | 0.24 ± 0.01 | 0.29 ± 0.02<br>§ |
| | | STR/Ort | 0.22 ± 0.01 | 0.22 ± 0.01 | 0.27 ± 0.02 | 0.24 ± 0.01 | 0.22 ± 0.02 | 0.3 ± 0.01<br>§§ |

These compartmentalised trends were mostly retained in old mice; extracondylar trabeculae were less stiff than cortical bone (both p=0.002) in CBAs and STR/Orts, and the predicted modulus in trabecular bone was lower across both condyles when compared to that in CBAs (lateral, p=0.0003; medial, p=0.007, Table 1). While age-related reductions in predicted modulus were found only in medial condylar trabecular bone in CBA (p=0.02), ageing STR/Orts were characterised by significantly stiffer lateral and medial SCP (p=0.0002 and p=0.0005) while neighbouring trabeculae (lateral, p=0.006 and medial, p=0.01) and extracondylar ROIs had lower predicted moduli in comparison to 10-week-old mice. We also calculated predicted Poisson’s ratio, which was largely uniform across all ROIs in 10- and 40-week-old CBA and 40-week-old STR/Ort epiphyses, with a modest reduction restricted to the lateral SCP (vs. medial, p=0.04). These data indicate that OA arising in the STR/Ort mouse involves regionalised and asymmetrical alterations in epiphyseal stiffness from 10 weeks of age, which arise without changes in volumetric compliance or in Poisson’s ratio, pinpointing a loss of load bearing capacity rather than mechanical performance as the primary deficit in incipient OA joint degeneration.

### Epiphyseal microarchitecture predicts susceptibility to load-induced deformation of the subchondral plate in advance of OA

To explore whether regional and asymmetrical divergence in epiphyseal strain distribution may arise due to incongruous skeletal microarchitecture, we next performed gross and microstructural evaluation of CBA and STR/Ort epiphyses (Fig. 4A and Supplementary Table S3). At 10-weeks of age, STR/Ort mice possessed significantly thicker SCP with greater volume on the medial tibial condyle (most OA-affected, Supplementary Fig. S2B) (p=0.0002 and p=0.0023, respectively) while the lateral condyle resembled that of healthy CBA mice. Significant asymmetry in the SCP was also evident in CBA mice albeit to a lesser extent wherein the medial SCP was thicker (p<0.0001) but with a lower volume (p=0.04) than the lateral SCP. This condylar asymmetry was also observed in STR/Orts; thicker yet volumetrically smaller trabeculae were present medially (p=0.001 and p=0.002, respectively). While trabecular thickness between condyles was comparable in CBA joint of the same age, trabecular volume was comparatively lower medially (p=0.0002). In extracondylar regions, cortices were thicker in STR/Orts than CBAs (p=0.004) with extracondylar trabecular bone being thinner across both genotypes (p<0.0001).

**Figure 4.**
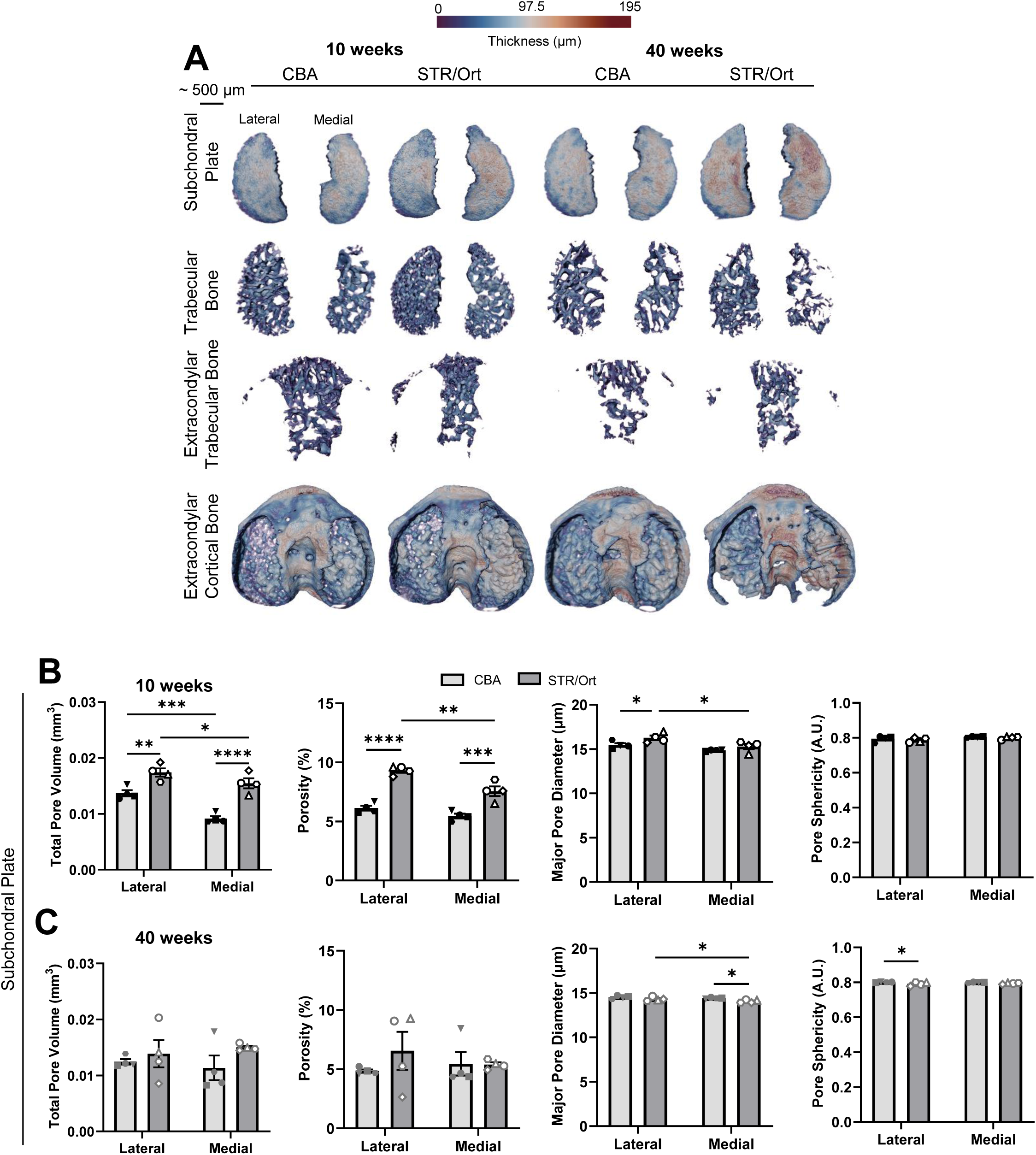
STR/Ort tibiae exhibit microstructural alterations in the subchondral plate that predispose to age-related osteoarthritis. Volumetric thickness maps of SCP, trabecular bone from the lateral and medial condyles of 10 week and 40-week-old CBA and STR/Ort epiphyses in addition to ECTB and extracondylar cortical bone ECC (A). Quantification of total pore volume, porosity, major pore diameter and pore sphericity reveals presence of greater pore volume and porosity in STR/Orts at 10-weeks of age (B) is lost with age (C) wherein STR/Ort porosity is comparable to CBAs. Data are presented as mean ± SEM with symbols representing individual animals (N=4/age/genotype). Statistical significance between condyles was assessed using linear mixed model analyses with Fisher’s LSD post-hoc test (C and D; *p<0.05, **p<0.01).

By 40 weeks of age, medial SCP thickening was still evident in CBA (p=0.005) and STR/Ort (p=0.008) epiphyses, and this only became more pronounced with age (both p=0.02). Ageing in both strains was accompanied by a loss of trabecular volume which was evident in the lateral condyle of CBA (p=0.001) and in across both condyles in STR/Ort epiphyses (p=0.004 and p=0.03, respectively). Prominent extracondylar corticalisation was evident in both CBA (thickness: p<0.0001 and volume: p=0.0009) and STR/Ort epiphyses (thickness: p=0.007 and volume: p=0.008) which occurred independently of further age-related changes to the extracondylar trabeculae bone. These data together suggest that both healthy and older OA-prone mice are characterised by SCP thickening and corticalisation which is unlikely to contribute to the greater load-induced deformation generated in STR/Ort epiphyses.

Intriguingly, total SCP pore volume was significantly greater in 10-week-old STR/Ort than age-matched CBA across both condyles (lateral: p=0.002 and medial: p<0.0001, Fig. 5B), which was linked to complementary genotype differences in percentage porosity (lateral: p<0.0001 and medial: p=0.0001). Within this, the lateral SCP exhibited greater total pore volume than the medial condyle (CBA, p=0.0001 and STR/Ort, p=0.01) while percentage porosity was exclusively higher laterally in STR/Ort epiphyses (p=0.003). Further interrogation revealed that these laterally-located SCP pores in STR/Orts had larger major pore diameters than those located within the medial condyle (p=0.01) and were additionally wider than those in the lateral CBA SCP (p=0.03) while pore sphericity remained consistent across animals and condyles. By 40 weeks of age, the differences in total pore volume and percentage porosity were no longer apparent (Fig. 5C). Instead, subtle reductions in major pore diameter were detected in the medial SCP of STR/Ort epiphyses (p=0.01). In this region, CBAs had greater pore diameter (p=0.02) while pores in the lateral SCP were more spherical than those in STR/Ort joints. These data indicate that OA-prone epiphyses exhibit regional and asymmetrical divergence, with greater bone quantity but more porous SCP, which align with raised epiphyseal strains, suggesting that the latter may arise due to incompatible alterations to skeletal microarchitecture.

**Figure 5.**
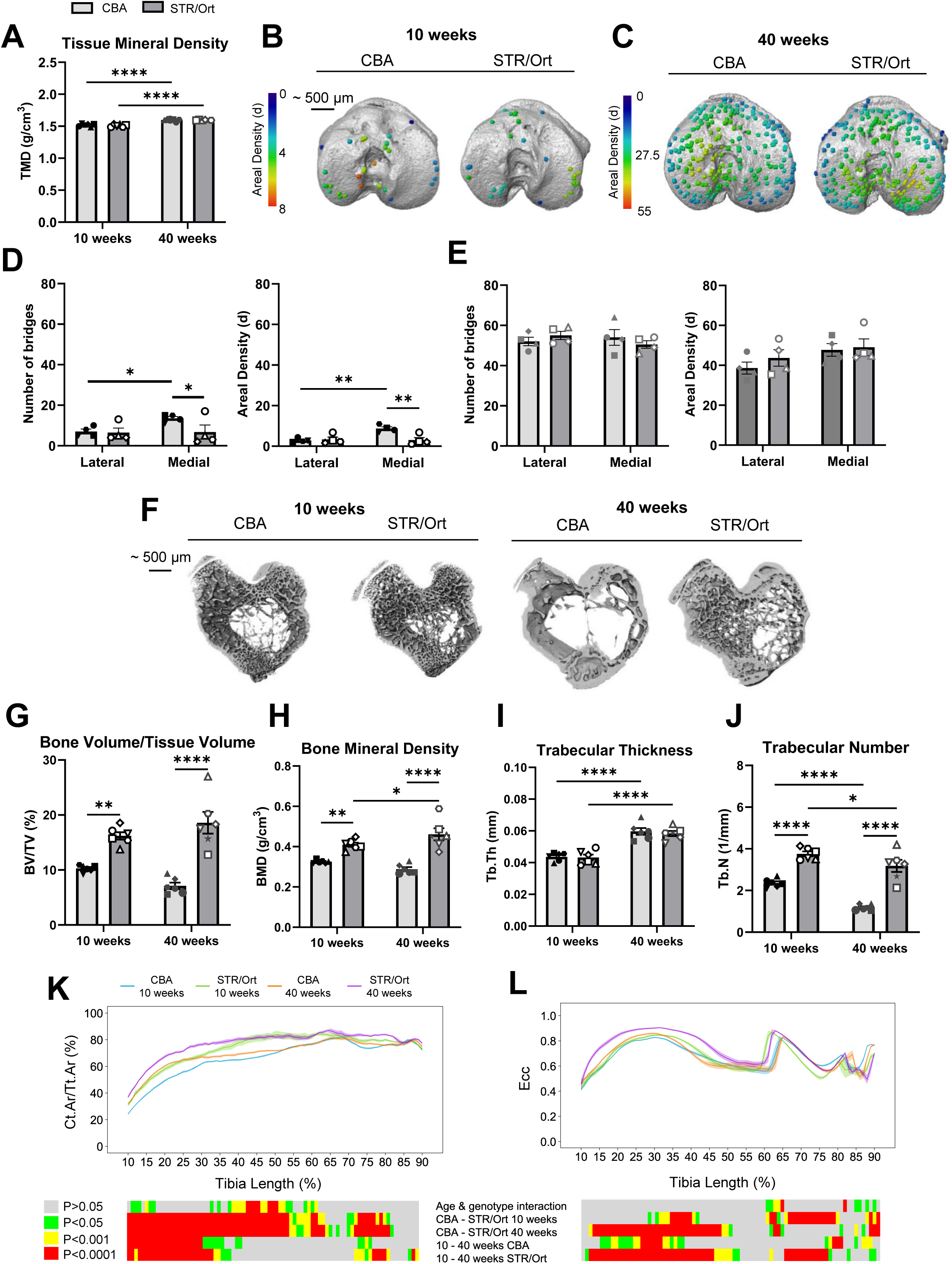
Enhanced metaphyseal, but not epiphyseal, bone mass is accompanied by proximal curvature in STR/Ort epiphyses. MicroCT-based tibial morphometric analyses reveal comparable epiphyseal TMD (A) between CBAs and STR/Orts of the same age, which further increase with age. Examination of the location and areal densities of growth plate bridges in 10- and 40-week-old CBA and STR/Orts (B and C, respectively) reveal greater bridge number and areal density in the lateral epiphyses of CBA mice at 10 weeks of age which is lost with age by 40-weeks (D and E, respectively). Representative 3D rendering of reconstructed microCT images of the tibial metaphyseal trabecular bone of 10- and 40-week-old CBA and STR/Ort mice (F). Morphometric analyses of metaphyseal trabecular BV/TV (G), BMD (H), Tb.Th (I) and Tb.N (J) indicate high bone mass of STR/Ort trabecular bone is maintained with age. Tibial Ct.Ar/Tt.Ar (K) and eccentricity (Ecc; J) were evaluated along the length of the tibia (10-90% of tibial length) are herein presented as line graphs representing mean ± SEM. Graphical heat maps beneath show statistical significance at spatially matched locations along the tibial length with age (10 versus 40 weeks) and between genotypes (CBA versus STR/Ort). Data are presented as mean ± SEM with symbols representing individual animals (N=4/age/genotype; D and E or N=6/age/genotype; A and G-L). Statistical significance between condyles, ages and genotypes was assessed using linear mixed model analyses with Fisher’s LSD post-hoc test (A, D-E and G-J, *p<0.05, **p<0.01, ***p<0.001 and ****p<0.0001) or two-way ANOVA with Tukey’s post-hoc test (K and L; green, p<0.05; yellow, p<0.001; and red, p<0.0001).

To explore links between strain-induced epiphyseal behaviours and metaphyseal/diaphyseal tibial architecture, we also performed microCT on whole tibiae and conducted morphometric analyses on the epiphysis, growth plate, trabecular and cortical bone compartments of 10 and 40-week-old CBA and STR/Ort. Tissue mineral density (TMD, Fig. 5A) increased with age (p<0.0001) but was not dissimilar in age-matched CBA and STR/Ort epiphyses, while epiphyseal volume (Supplementary Fig. S6A) was consistent in all groups. Growth plate bridge quantification revealed significantly greater bridge number and areal density on the medial than lateral condyle of 10-week-old CBA joints (p=0.05 and p=0.004, respectively) (Fig. 5B and D). In contrast, 10-week-old STR/Ort joints exhibited latero-medially matched bridge number and areal density which were lower medially than in CBA. Ageing resulted in significantly increased bridge number and density in both condyles in both CBA and STR/Ort mice (p<0.0001), which were comparable across all regions at 40 weeks (Fig. 5C and E).

Morphometric analyses of metaphyseal trabecular bone showed that 10-week STR/Ort mice had a high bone mass phenotype (Fig. 5F), characterised by greater bone volume/tissue volume (BV/TV; p=0.001, Fig. 5G), BMD (p=0.002, Fig. 5H) and trabecular number (Tb.N; p<0.0001) and trabecular thickness (Tb.Th; p<0.0001, Fig. 5G-J) than CBA mice. These differences were also prominent in 40-week-old animals (p<0.0001, Fig. 5I-J), with healthy ageing in CBAs characterised by increased Tb.Th (p<0.0001) and reduction in Tb.N (p<0.0001) while modest increases in BMD (p=0.04) and Tb.Th (p<0.0001) and reductions in Tb.N (p=0.02) characterise ageing in STR/Orts. Additional trabecular parameters that exhibited clear genotype and age-specific differences including trabecular spacing (Tb.Sp), trabecular pattern factor (Tb.Pf), degree of anisotropy, connectivity density (Conn.D) are shown in the Supplementary Information (Supplementary Fig. S6B-F). STR/Ort tibial cortices possessed greater cortical area/tissue area (Ct.Ar/Tt.Ar) and eccentricity changes than CBA tibiae at both ages (Fig. 5K and L) despite being shorter in length at 10-weeks of age (p=0.05, Supplementary Fig. S6G and H). Additional cortical shape metrics including minimum and maximum moments of inertia (*I_min_*, and *I_max_*, K, Supplementary Fig. S6I and J, respectively), resistance to torsion (*J*, L; Supplementary Fig. S6K) and cortical thickness (Ct.Th, Supplementary Fig. S6L) were solely altered with age. These analyses disclose prominent, spatially restricted tibial mass and geometry differences in 10-week-old STR/Ort that undergo distinct changes in ageing that align with divergent biomechanical performance of the epiphysis.

## Discussion

By integrating sCT and DVC we have redefined how loads are accommodated in intact healthy ageing joints and revealed that aberrant, regionally biased load-induced deformation arises in OA-prone joints prior to histological disease onset. In contrast to the uniform and symmetrically distributed strain fields generated in response to applied load in healthy CBA knees, elevated deformability and laterally-skewed load-induced compression and tension, signifying mechanical insufficiency, prefigured the initiation of OA in STR/Ort mice. Our use of regionalised DVC and FE-based analyses highlights that high strain foci arising within the SCP in STR/Ort joints are likely due to microarchitectural, but not material incongruities, implying that mechanical SCP vulnerability impairs the efficiency of load transmission seen in healthy CBA joints. Our data are consistent with the hypothesis that heightened microarchitectural porosity in the SCP permits inappropriate accumulation of load-induced strains within the epiphysis which predict emerging OA. Collectively, our spatially-correlated and multiscale approach has provided direct experimental evidence that nanoscale mechanical heterogeneity precedes overt OA pathology and can be used to define a preclinical mechanical signature of joint vulnerability.

Earlier non-invasive in vivo loading models have been instrumental in defining how skeletal elements adapt to axial mechanical load stimuli. In the tibial loading model of De Souza, Matsuura (37), delivery of 12 N dynamic loads induced periosteal expansion, cortical thickening and changes in metaphyseal trabecular architecture. Similarly, Poulet et al. (38) found that exposure to a single 9 N compressive load was sufficient to induce focal cartilage injury, while repeated load episodes produced progressive OA-like changes in the lateral femoral condyle, attributable to valgus joint positioning. While well suited to studying mechanoadaptative responses over periods of days to weeks, these approaches did not permit quantification of the instantaneous 3D strain fields that develop during loading. Our pioneering sCT-based in situ approaches (12) advanced this capability by coupling sCT imaging with DVC to show that OA-prone STR/Ort joints exhibit distinct chondrocyte lacunar organisation that is aligned with local indenter-induced strain accumulation and a failure to propagate strains from the articular surface to the subchondral bone. Building on these advances, we have herein tailored the non-surgical knee joint model of Poulet et al., (38) for sCT wherein in situ displacement-controlled loads could be delivered to the entire knee within an intact hindlimb and load transmission across the tibial epiphysis assessed by DVC. This involved the machining of identical loading cups to ensure physiological femur-tibia alignment while accommodating for load paths that reflect native physiologically oriented joint mechanics. In our approach, each displacement step was followed by a dedicated stress-relaxation period to allow force stabilisation while simultaneously mitigating for viscoelastic drift that would otherwise introduce motion artefacts and inherently compromise DVC accuracy. When combined with sCT, which enables visualisation of chondrocyte and osteocyte lacunae which constitute as image “texture”, DVC can be performed to compute full-field displacement and consequently strain (22). However, the stress-relaxation periods required in our protocol render these measurements quasi-static and therefore preclude the characterisation of strain-dependent behaviour of cartilage and surrounding soft tissues. Resolving such rapid viscoelastic behaviour would necessitate tomographic imaging at much faster, millisecond-scale acquisition rates. Nonetheless, these methodological refinements have enabled us to resolve regional heterogeneity in nanoscale strains across the epiphyses in physiologically loaded joints that age healthily, and in OA-prone joints, before and after disease onset, which were previously inaccessible to exploration using such load paradigms.

STR/Ort mice are amongst the most comprehensively characterised models of spontaneous OA, in which emergence invariably originates in the medial tibial plateau with signs of OA pathology in hyaline cartilage arising from 18-20 weeks (35, 36, 39–41). OA prevalence and development in males of this strain (34) is also marked by subchondral sclerosis and osteophytosis, closely mirroring primary OA in humans (36). Some early abnormalities have been described before histologically-defined cartilage damage, including a defect in endochondral ossification wherein accelerated long bone growth arises due to altered growth plate dynamics (15). Although several molecular pathways involving sclerostin, NF-ĸB signalling, members of the SIBLING family and matrix metalloproteinases have been implicated as early promoters of articular cartilage loss and subchondral sclerosis in STR/Ort (15, 35, 36), mounting evidence indicates that these molecular changes may be coupled to, and likely precipitated by, the fundamental mechanical compromise that is inherent to this strain. The notion that OA in STR/Orts is mechanically driven is indeed supported by observations of medial patella dislocation and anterior cruciate ligament weakening indicating inherent joint instability is linked with OA vulnerability (39). Like in many – likely all – forms of OA, cyclic mechanical loading is known to exacerbate medial tibial articular cartilage loss and subchondral bone thickening in STR/Ort knee joints, further linking inherently unstable joint mechanics to OA vulnerability (35, 42). In contrast, although focal cartilage lesions can be induced in the lateral femur of CBA knee joints in response to traumatic in vivo mechanical loading (35, 36), these mice exhibit very low, if any, susceptibility to OA. It is tempting to speculate that our data indicate that greater compressive deformability of the STR/Ort epiphyses precedes and may underpin their increased vulnerability to OA onset.

Our full-field DVC analyses build substantially upon our prior work (12) which used indenters to show that strain localisation in response to in situ loading occurred in the subchondral bone of CBA, but instead translocated to the calcified cartilage in STR/Ort joints. While this study served to focus attention upon these osteochondral interfaces, our approach herein combining whole epiphyseal DVC with regionalised anatomical subdivision, unequivocally demonstrates, for the first time that abnormalities in focal strain concentration at emerging OA stages in STR/Ort joints are spatially compartmentalised and exhibit marked condylar asymmetry. In extension, we also found that physiological load-induced deformation is uniformly and symmetrically dissipated through the SCP plate and underlying trabecular network to engender the highest tensile and compressive strains in the proximal growth plate, a site far-removed from the articulating joint regions. In stark contrast, the substantially higher magnitudes experienced within lateral SCP of STR/Ort joints under identical load, is consistent with the lateral-posterior joint loading known to be evoked in this model (38) and supports the concept that mechanical asymmetry, particularly in the SCP is a strong determinant of OA susceptibility.

It is therefore of major significance that our DVC findings and FE modelling also show that these latero-medial asymmetries are seemingly not explained by distinct material properties in these regions but are instead aligned with regional differences in SCP porosity. This implies that loss of load bearing capacity rather than mechanical performance (evaluated through characterisation of material properties) is the primary deficit that emerges in spontaneous OA that develops in STR/Ort mice. In agreement with our DVC findings, FE modelling showed comparable stiffness across lateral and medial tibial condyles while STR/Orts joints harbour significantly reduced SCP modulus with the most pronounced deficit observed laterally. This reduction in stiffness directly aligns with elevated and spatially broadened strain fields, confirming that these joints lack the load-bearing capacity required to effectively redistribute and transmit physiological loads. Although trabecular stiffness was also lower in STR/Orts, the lateral strain bias was not evident in this compartment highlighting that SCP insufficiency, rather than alterations in trabecular material properties is likely to be the primary driver of aberrant load transmission in these OA-prone joints.

We found that Poisson’s ratio was largely similar across all regions and ages, demonstrating that while intrinsic deformability is preserved, isolated reductions in stiffness coupled with thicker extracondylar cortical bone, or epiphyseal corticalisation, does not predispose STR/Ort epiphyses to early mechanical failure. Our FE-derived insights indeed refine and extend the conclusions of Poulet et al., (43) whose modelling approach disclosed that unusually thick articular cartilage evident at all ages in STR/Ort mice, effectively attenuates contact stresses at the joint surface to confer protection against load-induced lesions that readily develop in CBA mice. While previous studies have provided evidence for lower articular cartilage stiffness in the medial condyle of STR/Ort joints (44), our data show that the principal mechanical deficit instead lies within the SCP.

Our observations of altered growth plate bridging provide a parallel structural explanation for the strain patterns and align with previous descriptions of abnormal epiphyseal fusion in STR/Ort mice (15). In CBA mice, these bridges were enriched medially at 10 weeks of age, indicating that epiphyseal fusion progresses in a regionalised manner until bridge distribution is latero-medially balanced by 40-weeks of age. In young STR/Ort, however, we found fewer and more symmetrically distributed bridges which may reflect a loss of the normal spatial coupling between local mechanics and endochondral maturation. By 40-weeks of age, bridge number and areal densities were spatially conserved in both mouse genotypes, suggesting that bridge formation may arise via altered endochondral processes that bypass the requirement for mechanical guidance that characterises long bone growth in CBA mice. In our previous work (45) we showed that individual bridges were capable of acting as stress concentrators to enable transfer of epiphyseal strain across the growth plate to the proximal metaphysis, and that repetitive loading can actively modulate bridge formation (46), consistent with long standing evidence that the growth plate is acutely load sensitive (47). Importantly, our findings resonate with the model proposed by Xie et al., (48) in which the secondary ossification centre, which developmentally defines the epiphysis, functions to shield the mechanically vulnerable hypertrophic growth plate chondrocytes from excessive stress; implying that any inherent disturbance to growth plate bridging may generate compromised load redistribution, thus reinforcing the emergence of the raised focal strain concentrations that we have resolved herein.

Marked architectural divergence between CBA and STR/Ort tibiae provide promising insight into the link between the structural and mechanical basis for the impaired epiphyseal load dissipation observed by imaging intact, in situ loaded fresh murine knee joints. Early medial SCP and trabecular thickening that would typically enhance the mechanical strength of the epiphysis is offset by elevations in lateral SCP porosity and total pore volume in STR/Ort joints, even prior to OA. STR/Ort mice exhibited a high-bone mass phenotype in metaphyseal/diaphyseal bone which is consistent with earlier observations of Pasold et al., (49) and (50). Here, the metaphyseal high bone mass phenotype is accompanied by diaphyseal geometric abnormalities and regionalised corticalisation, indicative of enhanced regionalised stiffness which may potentiate the unfavourable translation of axial loads towards the epiphysis. Collectively these data support a model in which mis-patterning of long bone microarchitecture in STR/Orts during early life leads to the disturbance to epiphyseal structure, consequently leading to alterations in habitual load paths. Together these interpretations are in support of a unified view that OA susceptibility in STR/Orts arises not from isolated defects in any one area of the tibia, but rather from a collective failure across regions of the long bone. As a result, adoption of this architecture across scales is primed to generate high magnitude, laterally biased epiphyseal strains that precede overt OA.

Together, our sCT-DVC approach resolves nanoscale epiphyseal in situ load transfer and exposes preclinical mechanical vulnerabilities that precede overt onset of OA in articular cartilage. Regionalised DVC-validated FE modelling and anatomical profiling uncovers mechanobiological aberrations in which reduced and laterally biased SCP stiffness and elevated porosity within the epiphysis, combined with altered long bone geometry and gross architecture, induces focal strain amplification in the epiphysis long before overt cartilage degeneration. Herein we have identified a biomechanical fingerprint, comprising regional SCP modulus, SCP pore architecture and growth plate-adjacent strain gradients that can serve as powerful early marker for stratifying joint vulnerability. Although synchrotron-based in situ mechanical testing inevitably limits cohort size, this approach has yielded an unparalleled discovery platform for examining how genetic, developmental or therapeutic perturbations influence load transfer across joint interfaces. Beyond OA, these methods open new opportunities to interrogate multiscale mechanobiology across the skeletal system and can be used to evaluate the efficacy of interventions intended to normalise joint mechanics, while defining how nanoscale mechanical environments evolve following surgical, regenerative or prosthetic reconstruction. Thus, our work establishes a scalable mechanistic framework for identifying mechanical joint failure in OA and is poised to accelerate biomarker development and shorten the translational pathway for disease-modifying OA therapies.

## Methods

### Animals and sample preparation

All experimental procedures were performed in accordance with the UK Animals (Scientific Procedures) Act of 1986, UK Home Office Regulations and Royal Veterinary College, UK institutional guidelines. Male CBA (Charles River, UK) and STR/Ort (in-house) mice were kept in polypropylene cages at 21 ± 2 °C and subjected to 12hr:12hr light/dark cycles. All mice were fed a standard RM1 maintenance diet (No.1; Special Diet Services, Witham, UK) and water ad libitum. Male STR/Ort mice, maintained through brother-sister pairing, were examined at ages prior to the onset of OA (8-11 weeks) and at advanced OA stages (40 weeks) (36) and were compared to age-matched male CBA mice (healthy parental control strain; (35, 39) (N=6/age group). Mice were euthanised by cervical dislocation, after which whole left hindlimbs were removed and skinned leaving all soft tissues and musculature intact. Individual limbs were wrapped in 1X phosphate buffered (PBS) saline-soaked gauze and frozen at –80 °C in sealed polypropylene tubes until in situ testing and sCT. After this, whole hindlimbs were fixed in 4% neutral buffered formalin (v/v) for 48 hours at 4 °C then stored in 70% ethanol (v/v) until further analysis.

### In situ loading

Hindlimbs were thawed at room temperature prior to mounting in a Deben CT500 in situ compression stage (Deben UK LTD, UK) equipped with a 100 N loadcell and a custom-designed open frame comprising an aluminium alloy crossbar (grade 6082, Atomic Precision, UK) bridging two carbon fibre epoxy rods (Ø 6 mm ± 0.2mm and 120 mm length; Carbon Fibre Profiles, UK) threaded into an aluminium alloy base cover (grade 6082, Atomic Precision, UK). This open frame configuration facilitates easy access for precise sample mounting and alignment procedures while enhancing X-ray transmission efficiency and maximising signal intensity at the detector. Additionally incorporated within the Deben CT500 rig was an actuator with a linear encoder that provided 10 mm compression range with a displacement resolution of 3 μm and ± 1% accuracy across the full measurement range. A pair of vertically aligned bespoke sample holders (upper cup supporting the flexed knee joint and lower cup for securing the ankle flexed at 45 °; (37), machined in polyether ether ketone (Atomic Precision, UK), were used to mount whole hindlimbs for in situ mechanical experiments. Hindlimbs were subjected to axial compression, applied in the z-direction upwards from the lower cup, prior to the acquisition of sCT images. A baseline 2 N load was primarily applied to secure the knee and ankle in deep flexion (38, 43) with the first sCT image acquired following a 15-minute period given to allow for stress relaxation to minimise motion artefacts. Three successive sCT scans, also taken following a 15-minute stress-relaxation period, were acquired upon incremental displacement of 20 µm, 70 µm and 170 µm using a motor speed of 0.1 mm/min.

### Phase contrast sCT

High-resolution sCT images of in situ loaded murine knees were acquired using a filtered white X-ray beam (2.33 mm aluminium and 0.41 mm copper) with an average energy of 63 keV, on the BM05 beamline at the European Synchrotron Radiation Facility, Grenoble, France, at an isotropic voxel size of 1.45 µm. Incoming filtered X-rays were magnified with a 5x objective (NA=0.21) and detected using a custom made 50 µm LuAG:Ce scintillator (Crytur, Czechia) combined with a PCO.edge 4.2 camera (2048 x 2048 pixels, PCO Imaging, Germany). Following the acquisition of 41 flatfield and 40 darkfield images, 6000 projections were acquired in half-acquisition fly-scan mode (51), with the centre of rotation positioned on the right edge of the image resulting in a final field of view of 5.21 mm x 2.97 mm. Projections were captured over a continuous 360° rotation with an angular step size of 0.06° and exposure time of 18 milliseconds. The radiation dose on the samples was estimated to be approximately 7.2 kGy per scan (28.4 kGy per sample). The propagation distance between the sample and detector was iteratively tested before being set to 0.25 m enabling optimum in-line phase contrast in sCT images.

Raw sCT projections were flatfield and darkfield corrected with greyscale intensities normalised to the synchrotron current (200 mA mode) ahead of tomographic reconstruction into 16-bit tiff stacks using the in-house PyHST2 software (version 2023a) (52). Following normalisation, Paganin phase-retrieval (δ/β=500) (53) was applied, along with a 2D unsharp mask (σ=1, coefficient=3.5) in preparation for filtered back-projection (54) and ring artefact removal (55).

### sCT image preprocessing prior to digital volume correlation

Following the tomographic reconstruction, full sCT datasets were imported into Avizo3D (version 2022.2, Thermo Fisher Scientific, USA) and prepared for the global approach DVC and anatomical segmentation. Images were cropped to the smallest possible size for the isolation of the knee joint, with the same crop dimensions applied to successive sCT images of the same knee joint acquired following displacement-induced deformation. Tibiae, restricted to epiphysis and metaphyses, were segmented in cropped sCT images acquired after the application of the 2 N load using a 3D region growing algorithm. Osteocyte and chondrocyte lacunae were filled using the “fill volume” function followed by a 3-pixel dilation and erosion step prior to the manual removal of the metaphyseal growth plate resulting in a binary label image of the tibial epiphysis with growth plate bridges. Surfaces were created from binarised tibial epiphysis segmentations (smoothing factor = 5) and were further simplified through modifying the distance between nodes prior to unstructured finite element tetrahedral mesh generation (31). Evaluation of DVC performance was determined through uncertainty evaluation, in which a pair of repeat reference images were acquired sequentially from a single knee joint without the addition of further load. A minimum distance of 30 voxels (43.5 µm) between nodes yielded an optimal displacement accuracy of 0.012 voxels and precision of 78.3 nm across the tibial epiphysis. This subsequently resulted in a mean strain accuracy of 53 µstrain and precision of 221 µstrain (see Supplementary Information for further details). Based on this, this mesh size was therefore deemed suitable to track deformation around osteocyte lacunae (diameter ∼ 5-15 µm, (56)) and chondrocyte lacunae (diameter ∼ 25–30 µm, (57)) and were created for each sample in successive DVC analyses which had an average number of 14229 ± 173 nodes corresponding to 53412 ± 997 tetrahedra.

### Bulk epiphyseal strain analysis

Prior to the execution of the DVC algorithm (XDVC module), cropped sCT images of deformed knee joints (N=4 per age/genotype) taken after the application of 20 µm, 70 µm, and 170 µm displacement were rigidly registered (correcting for translation and rotation) to baseline images of the same sample acquired under a 2 N load. FE tetrahedral meshes created from tibial epiphysis segmentations were applied to reference and deformed image datasets post-registration. Displacement fields were computed incrementally through correlation of the reference images with each successive displacement step for each knee joint: 2 N vs 20 µm (+20 µm), 2 N vs 70 µm (+70 µm), and 2 N vs 170 µm (+170 µm). The FE-based DVC employed herein enabled the estimation of a continuous 3D displacement field using a global optimisation procedure in which the correlation between registered reference and deformed image volumes is maximised while difference in greyscale values is minimised in least-squares approach. Elemental Green-Lagrange strain tensors were subsequently computed at each nodal location from the computed displacement fields upon convergence of the DVC algorithm. Point clouds were then extracted from the nodes of the DVC finite element meshes from stepwise correlations and thresholded to exclude strains above 10 000 µstrain (1%) in line with published literature (58–61) with subsequent strain distribution analyses performed on 11964 ± 602 nodes.

### Anatomical epiphyseal characterisation

Total cortical and trabecular compartments of tibial epiphyses from individual mice were semi-automatically segmented in cropped, reconstructed sCT images following the application of the 2 N baseline load using the method described by (62) in Avizo 3D. Minor adjustments in the approach were iteratively optimised in order to apply the published method to our images with smaller voxel size, these included; 1) removal of a non-local means filter as Paganin phase-retrieval was performed during tomographic reconstruction; and 2) ball erosion/dilation operations were increased to 9 pixels and ball-closing operation was increased to 75 pixels. Briefly, epiphyseal cortices were segmented to isolate the extracondylar cortical bone from the subchondral plate. Medial and lateral subchondral plate segmentations were performed manually in the coronal plane in 29 µm increments and interpolated with all intermediate slices visually checked for interpolation errors and manually corrected when necessary. Final subchondral plate segmentations were subtracted from the total cortex, using the “arithmetic” function, to generate segmentations of the extracondylar cortex. To isolate the pores of the subchondral plate, the corresponding segmentations were eroded by 2 pixels then manually thresholded to isolate non-mineralised constituent tissues before a single-pixel ball erosion and dilation was applied to reduce noise prior to subsequent analysis. Extracted pores were exported as binary .tiff image stacks and subjected to 3D individual object analysis in CTAn (version 1.23.0.2 +; Bruker, Belgium) yielding measurements including volume, major diameter, and sphericity of individual pores from individual knee joints per tibial condyle. Pores smaller than 20 µm^3^ were assumed to be noise, those sized between 20-200 µm^3^ were assumed to be osteocyte lacunae while those sized between 200-2000 µm^3^ were assumed to be chondrocyte lacunae with vascular canals considered to be over 2000 µm^3^ in line with previously published literature (63–66) Trabecular bone was segmented next into lateral, medial and extracondylar compartments. Briefly, following overlay of the subchondral plate segmentations on 3D renders of the tibial epiphysis, the “lasso” tool was first used to draw around the edge of the lateral, then the medial SCP to isolate the total underlying lateral and medial condylar trabecular bone, respectively. The “arithmetic” function was then employed to subtract the lateral and medial condylar trabecular bone from the total trabecular bone segmentation resulting in the isolation of the extracondylar trabecular compartment.

To isolate DVC-derived strains from each anatomical compartment of the tibial epiphysis, volumetric bulk strains were imported into Avizo3D and spatially aligned and resampled to match the dimensions of the segmentation label generated above. The resampled volume subsequently underwent interactive thresholding to isolate the relevant anatomical structure, followed by sequential masking to remove background and refine the region of interest. The resultant masked volume, containing DVC-derived strains within anatomically defined regions of the tibial epiphysis, was subsequently volume rendered and used for further analysis.

### Spatial registration between DVC point cloud and FE model nodes

Before global optimisation of finite element-based modelling, DVC point clouds were registered to FE model nodes using a bespoke code in MATLAB (v. R2023b, MathWorks, Natick, US). A new DVC mesh (mesh 2) was generated with the same number of nodes and spatial positions before undergoing spatial alignment to the FE model mesh (Mesh 3). Principal strain values from the original DVC mesh (Mesh 1) were assigned to Mesh 2 using natural neighbour interpolation when nodes were within the Mesh 1 domain with further nearest-neighbour extrapolation performed for nodes falling outside of the Mesh 1 domain.

### Material property optimisation and FE analyses

FE models for CBA and STR/Ort tibial epiphyses were constructed based on the anatomical segmentations of the tibial epiphysis in reconstructed sCT images using Avizo3D of each tibial epiphysis. Briefly, binary segmentations from anatomical compartments were used to create surfaces as previously described. A convergence test was performed to identify optimal node spacing with each node assigned a homogenous elastic modulus of 150 MPa and Poisson’s ratio of 0.3. A convergence threshold of <1% strain change between successive mesh refinements was used to determine the node spacing of 25 voxels (36.3 µm) which was used for all subsequent analyses. The final converged meshes were exported in the .INP format for further optimisation (Supplementary Fig. 5A).

Model geometry, boundary conditions, and loading parameters were defined prior to global optimisation. Nodes located on the external surface of the epiphysis (referred to as Set-1) were fixed in all directions using encastre boundary conditions, while nodes on the inferior surface (designated as Set-2) were subjected to uniaxial compressive displacements of 20 μm, 70 μm, and 170 μm along the z-axis to replicate the in situ loading experiments. Finite element simulations were performed using Abaqus/Standard (version 2023, Simulia, Providence, USA), with the implicit solver used to compute the resulting strain fields within each anatomical region of the tibial epiphyses. To estimate the mechanical properties of each of the six anatomical compartments of the tibial epiphysis, an inverse optimisation approach was applied using a custom MATLAB pipeline (available at Zenodo, DOI: 10.5281/zenodo.15704567) in MATLAB). Within this framework, only the Elastic Modulus (10 MPa - 12,000 MPa) and Poisson’s ratio (0.1 - 0.45) were adjusted while geometrical boundary conditions and loading remained fixed. The resulting simulated first and third principal strain field were compared to spatially registered DVC-derived strain data with spatial correspondence evaluated following the calculation of the root-mean-square error (RMSE, Supplementary Fig. 5B and C). The MATLAB Global Optimisation Toolbox was used to iteratively adjust the material properties to achieve the global minimum RMSE and the final optimised set of regional material properties retained for further analysis.

### Bland–Altman plot

Bland–Altman plots were used to assess the agreement between the FE model with the optimal material properties and the DVC-derived point cloud strains. The first principal strain generated following 70 µm displacement was used for property optimisation while the first principal strain generated in response to 20 µm and 170 µm, along with the third principal strain generated following 20 µm, 70 µm, and 170 µm, were used for validation. The plots were generated by calculating the average and the difference between the FE modelled and DVC strain data in OriginPro (version 2023.b, Massachusetts, USA) (Supplementary Fig. S5D and E). The mean and standard deviation of the differences were then used to determine the 95% limits of agreement, defined as the mean ± 1.96 × SD.

### Micro-computed tomography

After in situ testing and sCT, left hindlimbs were imaged by micro-computed tomography (µCT) using a Skyscan 1176 scanner (Bruker, Belgium) (N=6/age/genotype). µCT scanning was performed at an isotropic voxel size of 4.98 µm, with the X-ray tube operating at 50 kV and 200 mA, using an exposure time of 960 ms with an 0.5 mm aluminium filter. Projections were collected every 0.6 ° over a continuous rotation of 180 ° and reconstructed into tomograms using NRecon (version 1.7.4.6; Bruker, Belgium). Additional µCT images of calcium hydroxyapatite phantoms with mineral densities of 0.25 g/cm^3^ and 0.75 g/cm^3^ (Bruker, Belgium) were acquired and reconstructed as above and used for attenuation coefficient calibration for the assessment of bone mineral density.

Prior to µCT-based morphometric analyses, µCT images were re-positioned in Dataviewer (version 1.5.6.2; Bruker, Belgium) to ensure consistent orientation and alignment for subsequent analyses. Tibiae were isolated by segmentation of reoriented µCT images in CTAn (version 1.23.0.2 +; Bruker, Belgium) prior to the measurement of tibial length. For 3D analyses of metaphyseal trabecular bone, a volume comprising 5% of the total bone length, beginning from the bottom of the growth plate where the primary spongiosa is no longer visible extending distally towards the diaphysis was used. Regions proximal to the unmineralized growth plate were used for 3D epiphyseal analyses. Datasets were segmented and binarised using a minimum threshold of 85 prior to the quantification of morphometric parameters (64, 67). For cortical bone analyses a 2D approach was conducted in CTAn following segmentation and binarisation using a minimum threshold of 80. Here, cortical bone was evaluated on a slice-by-slice basis thereby permitting the quantification of morphometric parameters along the tibial length. To ensure metaphyseal trabecular bone was not included in this analysis, 10% of the proximal and distal tibial length were excluded. 3D volume rendering of cortical and trabecular bone was achieved using Avizo3D.

Growth plate bridge analysis was performed as previously described by (45). Briefly, repositioned µCT datasets were imported into Avizo3D) and tibiae were segmented using a region-growing algorithm. The central regions of individual growth plate bridges across the width of the growth plate were identified in all image slices and planes (xy, xz and yz). The number and spatial distribution of growth plate bridges were determined using a bespoke in-house python script implemented in Avizo3D with growth plate bridges projected onto the tibial epiphysis and pseudo coloured to reflect the areal density at the bridge location.

### Histology and OARSI scoring

Following µCT, joints were decalcified in 10% ethylenediaminetetraacetic acid (EDTA) at 4 °C for 14 days before undergoing dehydration and processing for paraffin wax embedding, mounted in a frontal orientation, using standard procedures. Serial coronal sections of 6 µm thickness were cut throughout the entire knee joints. Every 4^th^ slide was selected for staining with Safranin-O (0.5% w/v in dH_2_0) and counterstained with Fast Green (0.2% w/v in dH20) staining (approximately 8-10 sections per joint) before imaging. Articular cartilage lesions were scored according to the internationally recognised OARSI scoring system (68). Scoring of cartilage lesioning across lateral and medial tibial condyles were conducted blindly by three independent scorers (AS, LAEE, LEB), with the maximum score and average score recorded for each sample and condyle. Agreement between scorers was deemed acceptable if the interclass correlation coefficient was 0.88, with a 95% confidence interval of 0.65 to 0.95.

### Statistical analysis

All investigators were blinded to the genotype and age of the animals throughout experimentation until data analysis. Statistical analyses were conducted using GraphPad Prism (version 10.5.0; Massachusetts, USA). Data normality was determined using the Shapiro-Wilk test. To assess the statistical differences in the distribution of DVC point cloud-derived strain between genotypes, the Kolmogorov-Smirnov test was used. One-way ANOVA followed by Dunnett’s post-hoc test was employed to evaluate the effect of increasing displacement on DVC-derived strain generated relative to the strain induced by +20 µm displacement. Linear mixed model analyses with Fisher’s LSD post-hoc test was used to assess condylar differences between the genotypes in each age group and for 3D trabecular morphometric analyses. For 2D analyses of cortical bone morphometry, Two-Way ANOVA test was performed to determine the interaction between genotype and age, using a bespoke code developed in R studio (version 2024.12.0, build 467; R Foundation for Statistical Computing, Austria), with statistical significance between region-matched locations presented in coloured heatmaps. Data are presented as mean ± standard error of the mean (SEM) and were considered statistically significant when p<0.05.

## Data availability

The full sCT data and FE models are available from the corresponding author upon reasonable request.

## Acknowledgements

Laboratory facilities were provided by the Royal Veterinary College, the University of Brighton and the Research Complex at Harwell. We thank C. Reinhardt and L. Sinclair (University of Manchester at Harwell) for access to the Deben CT500 mechanical testing rig and E. Burke O’Leary, C. Berruyer and V. Schoeppler for their assistance during beamtime at the European Synchrotron Radiation Facility (proposal LS-3124). This work was supported by the UKRI MRC (MR/V033506/1), the BLAST Network (BB/W01825X/1), Royal Academy of Engineering (CiET 1819/10) and the Chan Zuckerberg Initiative (CZIF2021-006424 and CZIF2022-316777).

## Author contributions

Conceptualisation: AS, LAEE, PDL, AAP, KAS; Methodology: AS, LAEE, LEB, KM, JC, ALP, PDL, AAP, KAS; Software: AS, LAEE, LEB, KM, JC, JB; Validation: AS, LAEE, LEB; Formal analysis: AS, LAEE, LEB, JC; Investigation: AS, LAEE, LEB, JC, ALP, JB; Resources: PDL, AAP and KAS; Data curation: AS, LAEE, LEB and JB; Writing – original draft: AS, LAEE, LEB, ALP, JC, PDL, AAP, KAS; Writing – review and editing: all authors; Visualisation: AS, LAEE, LEB, JC; Supervision: PDL, AAP and KAS; Project administration: PDL, AAP and KAS; Funding acquisition: PDL, AAP and KAS.

## References

1. Burr DB, Gallant MA. Bone remodelling in osteoarthritis. Nature reviews Rheumatology. 2012;8(11):665–73.

2. Hunter DJ, Bierma-Zeinstra S. Osteoarthritis. The Lancet. 2019;393(10182):1745–59.

3. Steinmetz JD, Culbreth GT, Haile LM, Rafferty Q, Lo J, Fukutaki KG, et al. Global, regional, and national burden of osteoarthritis, 1990&#x2013;2020 and projections to 2050: a systematic analysis for the Global Burden of Disease Study 2021. The Lancet Rheumatology. 2023;5(9):e508–e22.

4. Kolasinski SL, Neogi T, Hochberg MC, Oatis C, Guyatt G, Block J, et al. 2019 American College of Rheumatology/Arthritis Foundation Guideline for the Management of Osteoarthritis of the Hand, Hip, and Knee. Arthritis & rheumatology (Hoboken, NJ). 2020;72(2):220–33.

5. Oo WM, Hunter DJ. Disease modification in osteoarthritis: are we there yet? Clinical and experimental rheumatology. 2019;37(5):135–40.

6. Vincent TL, McClurg O, Troeberg L. The Extracellular Matrix of Articular Cartilage Controls the Bioavailability of Pericellular Matrix-Bound Growth Factors to Drive Tissue Homeostasis and Repair. Int J Mol Sci. 2022;23(11).

7. Pan J, Zhou X, Li W, Novotny JE, Doty SB, Wang L. In situ measurement of transport between subchondral bone and articular cartilage. J Orthop Res. 2009;27(10):1347–52.

8. Kamibayashi L, Wyss UP, Cooke TD, Zee B. Trabecular microstructure in the medial condyle of the proximal tibia of patients with knee osteoarthritis. Bone. 1995;17(1):27–35.

9. Hu YJ, Yu YE, Cooper HJ, Shah RP, Geller JA, Lu XL, et al. Mechanical and structural properties of articular cartilage and subchondral bone in human osteoarthritic knees. Journal of bone and mineral research : the official journal of the American Society for Bone and Mineral Research. 2024;39(8):1120–31.

10. Bobinac D, Spanjol J, Zoricic S, Maric I. Changes in articular cartilage and subchondral bone histomorphometry in osteoarthritic knee joints in humans. Bone. 2003;32(3):284–90.

11. Peters AE, Akhtar R, Comerford EJ, Bates KT. The effect of ageing and osteoarthritis on the mechanical properties of cartilage and bone in the human knee joint. Sci Rep. 2018;8(1):5931.

12. Madi K, Staines KA, Bay BK, Javaheri B, Geng H, Bodey AJ, et al. In situ characterization of nanoscale strains in loaded whole joints via synchrotron X-ray tomography. Nat Biomed Eng. 2020;4(3):343–54.

13. Li J, Yuan H, Wu M, Dong L, Zhang L, Shi H, et al. Quantitative assessment of murine articular cartilage and bone using X-ray phase-contrast imaging. PLoS One. 2014;9(11):e111939.

14. Xu H, Olivier C, Sajidy H, Pallu S, Portier H, Peyrin F, et al. Cell quantification at the osteochondral interface from synchrotron radiation phase contrast micro-computed tomography images using a deep learning approach. Sci Rep. 2024;14(1):29619.

15. Staines KA, Madi K, Mirczuk SM, Parker S, Burleigh A, Poulet B, et al. Endochondral Growth Defect and Deployment of Transient Chondrocyte Behaviors Underlie Osteoarthritis Onset in a Natural Murine Model. Arthritis & Rheumatology. 2016;68(4):880–91.

16. Marenzana M, Hagen CK, Borges PD, Endrizzi M, Szafraniec MB, Vincent TL, et al. Synchrotron-and laboratory-based X-ray phase-contrast imaging for imaging mouse articular cartilage in the absence of radiopaque contrast agents. Philos Trans A Math Phys Eng Sci. 2014;372(2010):20130127.

17. Li J, Zhong Z, Connor D, Mollenhauer J, Muehleman C. Phase-sensitive X-ray imaging of synovial joints. Osteoarthritis Cartilage. 2009;17(9):1193–6.

18. Honkanen MKM, Saukko AEA, Turunen MJ, Shaikh R, Prakash M, Lovric G, et al. Synchrotron MicroCT Reveals the Potential of the Dual Contrast Technique for Quantitative Assessment of Human Articular Cartilage Composition. J Orthop Res. 2020;38(3):563–73.

19. Li J, Zhong Z, Lidtke R, Kuettner KE, Peterfy C, Aliyeva E, et al. Radiography of soft tissue of the foot and ankle with diffraction enhanced imaging. Journal of anatomy. 2003;202(5):463–70.

20. Chappard C, Peyrin F, Bonnassie A, Lemineur G, Brunet-Imbault B, Lespessailles E, et al. Subchondral bone micro-architectural alterations in osteoarthritis: a synchrotron micro-computed tomography study. Osteoarthritis and Cartilage. 2006;14(3):215–23.

21. Fan X, Lee KM, Jones MWM, Howard D, Sun AR, Crawford R, et al. Spatial distribution of elements during osteoarthritis disease progression using synchrotron X-ray fluorescence microscopy. Sci Rep. 2023;13(1):10200.

22. Bay BK, Smith TS, Fyhrie DP, Saad M. Digital volume correlation: Three-dimensional strain mapping using X-ray tomography. Experimental Mechanics. 1999;39(3):217–26.

23. Palanca M, Cristofolini L, Dall’Ara E, Curto M, Innocente F, Danesi V, et al. Digital volume correlation can be used to estimate local strains in natural and augmented vertebrae: An organ-level study. Journal of Biomechanics. 2016;49(16):3882–90.

24. Dall’Ara E, Peña-Fernández M, Palanca M, Giorgi M, Cristofolini L, Tozzi G. Precision of Digital Volume Correlation Approaches for Strain Analysis in Bone Imaged with Micro-Computed Tomography at Different Dimensional Levels. Frontiers in Materials. 2017;Volume 4 - 2017.

25. Oliviero S, Giorgi M, Dall’Ara E. Validation of finite element models of the mouse tibia using digital volume correlation. Journal of the Mechanical Behavior of Biomedical Materials. 2018;86:172–84.

26. Giorgi M, Dall’Ara E. Variability in strain distribution in the mice tibia loading model: A preliminary study using digital volume correlation. Medical Engineering & Physics. 2018;62:7–16.

27. Davis S, Karali K, Zekonyte J, Roldo M, Blunn G. 3D full-field strain distribution across the osteochondral unit during osteoarthritis progression. VIEW. 2025;n/a(n/a):20250062.

28. Parmenter AL, Newham E, Sharma A, Disney CM, Deyhle H, Bosi F, et al. Variations in mineral prestrain, nanostructure, and microarchitecture play a role in intervertebral disc loading. Cell Biomaterials. 2025:100151.

29. Disney CM, Eckersley A, McConnell JC, Geng H, Bodey AJ, Hoyland JA, et al. Synchrotron tomography of intervertebral disc deformation quantified by digital volume correlation reveals microstructural influence on strain patterns. Acta biomaterialia. 2019;92:290–304.

30. Davis S, Karali A, Zekonyte J, Roldo M, Blunn G. Development of a method to investigate strain distribution across the cartilage-bone interface in guinea pig model of spontaneous osteoarthritis using lab-based contrast enhanced X-ray-computed tomography and digital volume correlation. Journal of the Mechanical Behavior of Biomedical Materials. 2023;144:105999.

31. Madi K, Tozzi G, Zhang Q, Tong J, Cossey A, Au A, et al. Computation of full-field displacements in a scaffold implant using digital volume correlation and finite element analysis. Medical engineering & physics. 2013;35(9):1298–312.

32. Kusins J, Knowles N, Columbus M, Oliviero S, Dall’Ara E, Athwal GS, et al. The Application of Digital Volume Correlation (DVC) to Evaluate Strain Predictions Generated by Finite Element Models of the Osteoarthritic Humeral Head. Ann Biomed Eng. 2020;48(12):2859–69.

33. Peña Fernández M, Sasso SJ, McPhee S, Black C, Kanczler J, Tozzi G, et al. Nonlinear micro finite element models based on digital volume correlation measurements predict early microdamage in newly formed bone. Journal of the Mechanical Behavior of Biomedical Materials. 2022;132:105303.

34. Walton M. Degenerative joint disease in the mouse knee; histological observations. The Journal of pathology. 1977;123(2):109–22.

35. Poulet B, Ulici V, Stone TC, Pead M, Gburcik V, Constantinou E, et al. Time-series transcriptional profiling yields new perspectives on susceptibility to murine osteoarthritis. Arthritis and rheumatism. 2012;64(10):3256–66.

36. Staines KA, Poulet B, Wentworth DN, Pitsillides AA. The STR/ort mouse model of spontaneous osteoarthritis - an update. Osteoarthritis Cartilage. 2017;25(6):802–8.

37. De Souza RL, Matsuura M, Eckstein F, Rawlinson SC, Lanyon LE, Pitsillides AA. Non-invasive axial loading of mouse tibiae increases cortical bone formation and modifies trabecular organization: a new model to study cortical and cancellous compartments in a single loaded element. Bone. 2005;37(6):810–8.

38. Poulet B, Hamilton RW, Shefelbine S, Pitsillides AA. Characterizing a novel and adjustable noninvasive murine joint loading model. Arthritis Rheum. 2011;63(1):137–47.

39. Walton M. Degenerative joint disease in the mouse knee; radiological and morphological observations. The Journal of pathology. 1977;123(2):97–107.

40. Chambers MG, Cox L, Chong L, Suri N, Cover P, Bayliss MT, et al. Matrix metalloproteinases and aggrecanases cleave aggrecan in different zones of normal cartilage but colocalize in the development of osteoarthritic lesions in STR/ort mice. Arthritis Rheum. 2001;44(6):1455–65.

41. Mason RM, Chambers MG, Flannelly J, Gaffen JD, Dudhia J, Bayliss MT. The STR/ort mouse and its use as a model of osteoarthritis. Osteoarthritis Cartilage. 2001;9(2):85–91.

42. Poulet B, de Souza R, Kent AV, Saxon L, Barker O, Wilson A, et al. Intermittent applied mechanical loading induces subchondral bone thickening that may be intensified locally by contiguous articular cartilage lesions. Osteoarthritis and Cartilage. 2015;23(6):940–8.

43. Poulet B, Westerhof TA, Hamilton RW, Shefelbine SJ, Pitsillides AA. Spontaneous osteoarthritis in Str/ort mice is unlikely due to greater vulnerability to mechanical trauma. Osteoarthritis Cartilage. 2013;21(5):756–63.

44. Lavoie JF, Sim S, Quenneville E, Garon M, Moreau A, Bushmann MD, et al. Mapping articular cartilage biomechanical properties of normal and osteoarthritis mice using indentation. Osteoarthritis and Cartilage. 2015;23:A254.

45. Staines KA, Madi K, Javaheri B, Lee PD, Pitsillides AA. A Computed Microtomography Method for Understanding Epiphyseal Growth Plate Fusion. Frontiers in Materials. 2018;Volume 4 - 2017.

46. Monzem S, Valkani D, Evans LAE, Chang YM, Pitsillides AA. Regional modular responses in different bone compartments to the anabolic effect of PTH (1-34) and axial loading in mice. Bone. 2023;170:116720.

47. Villemure I, Cloutier L, Matyas JR, Duncan NA. Non-uniform strain distribution within rat cartilaginous growth plate under uniaxial compression. Journal of Biomechanics. 2007;40(1):149–56.

48. Xie M, Gol’din P, Herdina AN, Estefa J, Medvedeva EV, Li L, et al. Secondary ossification center induces and protects growth plate structure. eLife. 2020;9:e55212.

49. Pasold J, Engelmann R, Keller J, Joost S, Marshall RP, Frerich B, et al. High bone mass in the STR/ort mouse results from increased bone formation and impaired bone resorption and is associated with extramedullary hematopoiesis. Journal of bone and mineral metabolism. 2013;31(1):71–81.

50. Javaheri B, Razi H, Piles M, de Souza R, Chang YM, Maric-Mur I, et al. Sexually dimorphic tibia shape is linked to natural osteoarthritis in STR/Ort mice. Osteoarthritis and Cartilage. 2018;26(6):807–17.

51. Kyrieleis A, Ibison M, Titarenko V, Withers PJ. Image stitching strategies for tomographic imaging of large objects at high resolution at synchrotron sources. Nuclear Instruments and Methods in Physics Research Section A: Accelerators, Spectrometers, Detectors and Associated Equipment. 2009;607(3):677–84.

52. Mirone A, Brun E, Gouillart E, Tafforeau P, Kieffer J. The PyHST2 hybrid distributed code for high speed tomographic reconstruction with iterative reconstruction and a priori knowledge capabilities. Nuclear Instruments and Methods in Physics Research Section B: Beam Interactions with Materials and Atoms. 2014;324:41–8.

53. Paganin D, Mayo SC, Gureyev TE, Miller PR, Wilkins SW. Simultaneous phase and amplitude extraction from a single defocused image of a homogeneous object. J Microsc. 2002;206(Pt 1):33–40.

54. van Aarle W, Palenstijn WJ, Cant J, Janssens E, Bleichrodt F, Dabravolski A, et al. Fast and flexible X-ray tomography using the ASTRA toolbox. Opt Express. 2016;24(22):25129–47.

55. Lyckegaard A, Johnson G, Tafforeau P. Correction of ring artifacts in X-ray tomographic images. International Journal of Tomography and Statistics. 2011;18:1–9.

56. Dallas SL, Prideaux M, Bonewald LF. The Osteocyte: An Endocrine Cell … and More. Endocrine Reviews. 2013;34(5):658–90.

57. Tang T, Zhong J, Hu J, Schemenz V, Davydok A, Brunner R, et al. Gradients in lacunar morphology and cartilage mineralization reflect the mechanical function of the mouse femoral head epiphysis. Acta biomaterialia. 2025;201:385–99.

58. Carriero A, Abela L, Pitsillides AA, Shefelbine SJ. Ex vivo determination of bone tissue strains for an in vivo mouse tibial loading model. J Biomech. 2014;47(10):2490–7.

59. Bayraktar HH, Morgan EF, Niebur GL, Morris GE, Wong EK, Keaveny TM. Comparison of the elastic and yield properties of human femoral trabecular and cortical bone tissue. J Biomech. 2004;37(1):27–35.

60. Oliviero S, Owen R, Reilly GC, Bellantuono I, Dall’Ara E. Optimization of the failure criterion in micro-Finite Element models of the mouse tibia for the non-invasive prediction of its failure load in preclinical applications. J Mech Behav Biomed Mater. 2021;113:104190.

61. Rubin CT, Lanyon LE. Dynamic strain similarity in vertebrates; an alternative to allometric limb bone scaling. Journal of Theoretical Biology. 1984;107(2):321–7.

62. Herbst EC, Felder AA, Evans LAE, Ajami S, Javaheri B, Pitsillides AA. A new straightforward method for semi-automated segmentation of trabecular bone from cortical bone in diverse and challenging morphologies. R Soc Open Sci. 2021;8(8):210408.

63. Mosey H, Núñez JA, Goring A, Clarkin CE, Staines KA, Lee PD, et al. Sost Deficiency does not Alter Bone’s Lacunar or Vascular Porosity in Mice. Frontiers in Materials. 2017;Volume 4 - 2017.

64. Javaheri B, Carriero A, Staines KA, Chang YM, Houston DA, Oldknow KJ, et al. Phospho1 deficiency transiently modifies bone architecture yet produces consistent modification in osteocyte differentiation and vascular porosity with ageing. Bone. 2015;81:277–91.

65. Carriero A, Doube M, Vogt M, Busse B, Zustin J, Levchuk A, et al. Altered lacunar and vascular porosity in osteogenesis imperfecta mouse bone as revealed by synchrotron tomography contributes to bone fragility. Bone. 2014;61:116–24.

66. Evans LAE. Imaging the articular osteochondral interface: University of London (Royal Veterinary College); 2023.

67. Javaheri B, Poulet B, Al-Jazzar AJ, de Souza R, Piles M, Hopkinson M, et al. Stable sulforaphane protects against gait anomalies and modifies bone microarchitecture in the spontaneous STR/Ort model of osteoarthritis. Bone. 2017;103:308–17.

68. Glasson SS, Chambers MG, Van Den Berg WB, Little CB. The OARSI histopathology initiative – recommendations for histological assessments of osteoarthritis in the mouse. Osteoarthritis and Cartilage. 2010;18:S17–S23.

