## Supplementary Information for "In situ profiling of nanoscale displacements uncovers mechano-architectural predictors of osteoarthritis emergence"

**DVC Uncertainty Measurement Results**

To determine the appropriate finite element tetrahedral mesh size for DVC analyses and for the evaluation of DVC performance, both systematic and random displacement error uncertainties and strain precision were determined using the DVC uncertainty algorithm in Avizo3D (XDVC module). Tibial epiphyses from repeat scans of the sample were segmented as described in the methods and used to generate a series of finite element tetrahedral meshes with varying minimum distance between nodes. A mesh size with a minimum node spacing of 30 voxels (43.5 µm) was identified as optimal, yielding mean displacements (referred to as accuracy) of 0.458 (x), 0.621 (y) and 0.0122 (z) voxels corresponding to 0.665, 0.901 and 0.018 µm (x, y and z, respectively). DVC precision, quantified as the standard deviation of the displacement field were 0.012 (x), 0.043 (y) and 0.054 (z) voxels which corresponded to 0.017, 0.063 and 0.079 µm (x, y and z, respectively). Similarly, average strain accuracy was 53 µstrain ($E_{xx}$ = 65 µstrain, $E_{yy}$ = 95 µstrain, $E_{zz}$ = 108 µstrain, $E_{xy}$ = 25 µstrain, $E_{yz}$ = 29 µstrain and $E_{xz}$ = 16 µstrain) while average precision was 221 µstrain ($E_{xx}$ = 233 µstrain, $E_{yy}$ = 181 µstrain, $E_{zz}$ = 415 µstrain, $E_{xy}$ = 127 µstrain, $E_{yz}$ = 178 µstrain and $E_{xz}$ = 196 µstrain).

**Supplementary Figures**

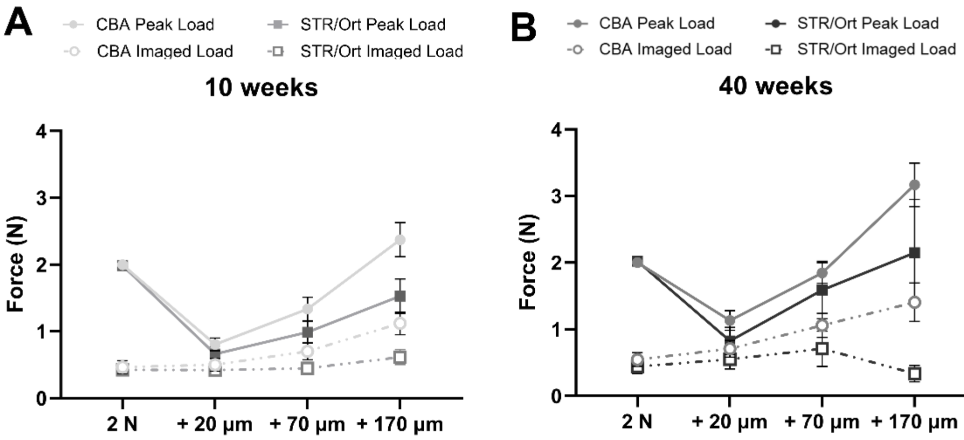


Supplementary Figure S1. Loading regime for in situ compression of 10- (A) and 40-week-old (B) CBA and STR/Ort knee joints. Following the application of a 2 N baseline load, knee joints were left for 15 minutes to permit stress-relaxation and sCT imaging was conducted following the stabilisation of the relaxed load henceforth referred to as the imaged load. Successive sCT scans were acquired at the imaged load in response to 20 µm, 70 µm, 170 µm displacement.

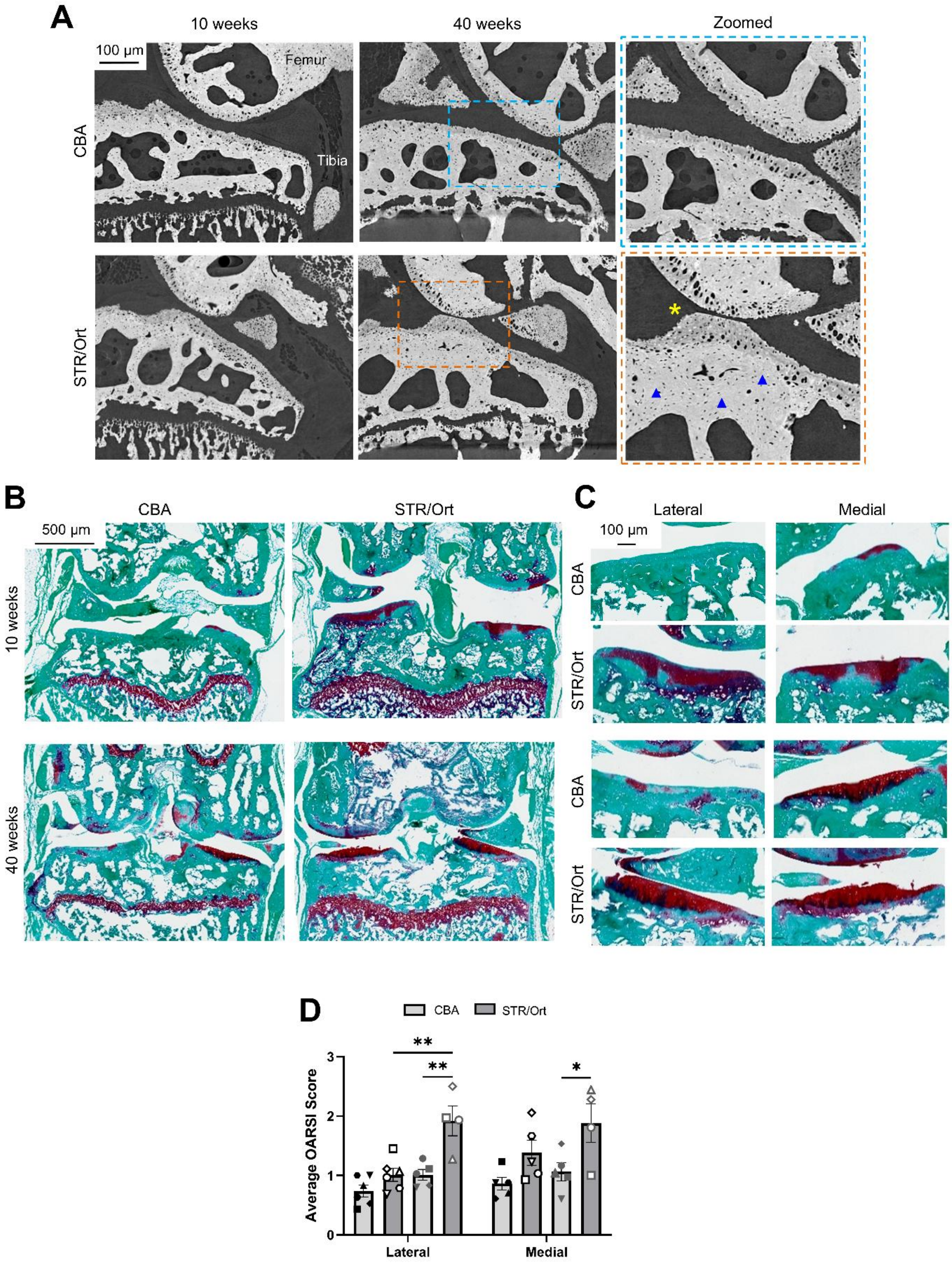


Supplementary Figure S2. Visualisation of epiphyseal microstructure sCT highlight osteoarthritis-associated pathology within the tibial epiphyses of STR/Ort knee joints with age (A). In CBAs, knee joint architecture is preserved with ageing (top panel), and are comparable to that in 10-week-old STR/Ort mice (bottom panel). By 40-weeks of age, advanced osteoarthritis is evident in zoomed images of STR/Ort knee joints (middle bottom, orange hatched box) and is characterised by joint space narrowing (yellow asterisk) and thickening of subchondral bone (blue arrowheads), which are not evident in the tibial epiphyses of age-matched CBA mice (blue hatched box). Histological examination following Safranin-O and Fast-Green staining of decalcified, paraffin-embedded knee joints post-sCT (B) highlights presence and location of cartilage lesions in the lateral and medial condyles of STR/Orts arise with age in zoomed images (C) and is reflected in greater average OARSI scores compared to CBAs (D). Data are presented as mean ± SEM with symbols representing individual animals (N=4/age/genotype). Statistical significance between condyles was assessed using linear mixed model analyses with Tukey's post-hoc test (D; *$p<0.05$, **$p<0.01$).

Supplementary Table S1. Median and interquartile range (IQR) of compressive strains in the tibial epiphyses of CBA and STR/Ort mice at 10- and 40-weeks of age following applied displacements of 20, 70 and 170 µm in situ. Data presented as average values for N=4 mice per age/genotype.

| | | | **CBA** | **STR/Ort** |
|---|---|---|---|---|
| **10 weeks** | 20 µm | **Median** | –0.0009 | -0.0012 |
| | | **IQR** | 0.0016 | 0.0017 |
| | 70 µm | **Median** | -0.0013 | -0.0014 |
| | | **IQR** | 0.0019 | 0.0022 |
| | 170 µm | **Median** | -0.0016 | -0.0017 |
| | | **IQR** | 0.0022 | 0.0023 |
| **40 weeks** | 20 µm | **Median** | –0.001 | -0.0013 |
| | | **IQR** | 0.0017 | 0.0019 |
| | 70 µm | **Median** | –0.0012 | -0.0013 |
| | | **IQR** | 0.0018 | 0.0019 |
| | 170 µm | **Median** | –0.0013 | -0.002 |
| | | **IQR** | 0.002 | 0.003 |

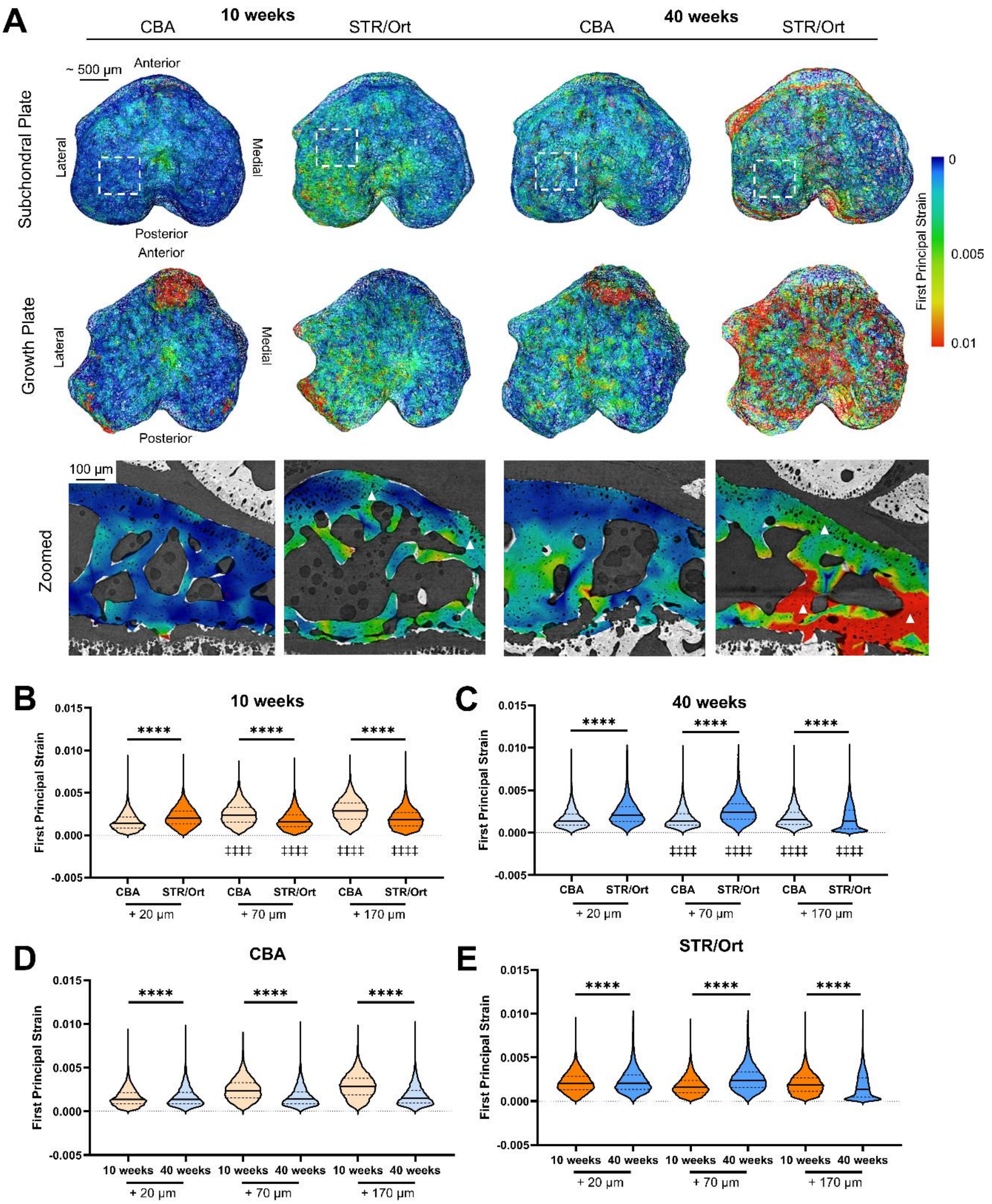


Supplementary Figure S3. DVC-computed tensile strains superimposed on finite element tetrahedral meshes of 10- and 40-week-old CBA and STR/Ort tibial epiphyses, viewed from the SCP (top row) and from the growth plate (bottom row) show localisation of high (red) and low (blue) first principal strain. In CBAs, tensile strains of high magnitude are confined to growth plate regions in 10- and 40-week-old animals. In STR/Orts, high tensile strains are evident across the SCP and growth plate and increase in magnitude with age. Zoomed view of tensile strain magnitude overlaid on sagittal sCT images show distal strain accumulation in CBA epiphyses in both ages, while in STR/Orts, higher magnitude strains are evident across the tibial epiphyses. Quantification of the distribution of tensile strains from point cloud nodes in response to displacement-induced loading in 10- (B) and 40-week-old (C) CBA and STR/Ort epiphyses are presented in violin plots with median value indicated by the solid line and upper and lower dashed lines representing 25$^{th}$ and 75$^{th}$ quartiles, respectively. Effect of age on tensile strain distribution in CBAs (D) and STR/Orts (E) in response to incremental displacement are shown. Individual violins represent DVC-derived strains from N=4/age/genotype. Statistical significance between CBAs and STR/Orts was assessed using the Kolmogorov-Smirnov test (**$p<0.01$, ***$p<0.001$ and ****$p<0.0001$). Statistical significance between displacement-induced load steps in CBA and STR/Ort mice relative to the strains induced in response to 20 μm displacement was assessed using One-way ANOVA with Dunnett's post-hoc test (‡‡‡‡$p<0.0001$).

Supplementary Table S2. Median and interquartile range (IQR) of tensile strains in the tibial epiphyses of CBA and STR/Ort mice at 10- and 40-weeks of age following applied displacements of 20, 70 and 170 µm in situ. Data presented as average values for N=4 mice/age/genotype.

| | | | CBA | STR/Ort |
|---|---|---|---|---|
| 10 weeks | 20 µm | **Median** | 0.0012 | 0.0019 |
| | | **IQR** | 0.0014 | 0.0021 |
| | 70 µm | **Median** | 0.0022 | 0.0015 |
| | | **IQR** | 0.0026 | 0.0017 |
| | 170 µm | **Median** | 0.0029 | 0.0017 |
| | | **IQR** | 0.0028 | 0.002 |
| 40 weeks | 20 µm | **Median** | 0.0011 | 0.0015 |
| | | **IQR** | 0.0019 | 0.0016 |
| | 70 µm | **Median** | 0.0012 | 0.0018 |
| | | **IQR** | 0.0016 | 0.0016 |
| | 170 µm | **Median** | 0.0013 | 0.0018 |
| | | **IQR** | 0.0025 | 0.0012 |

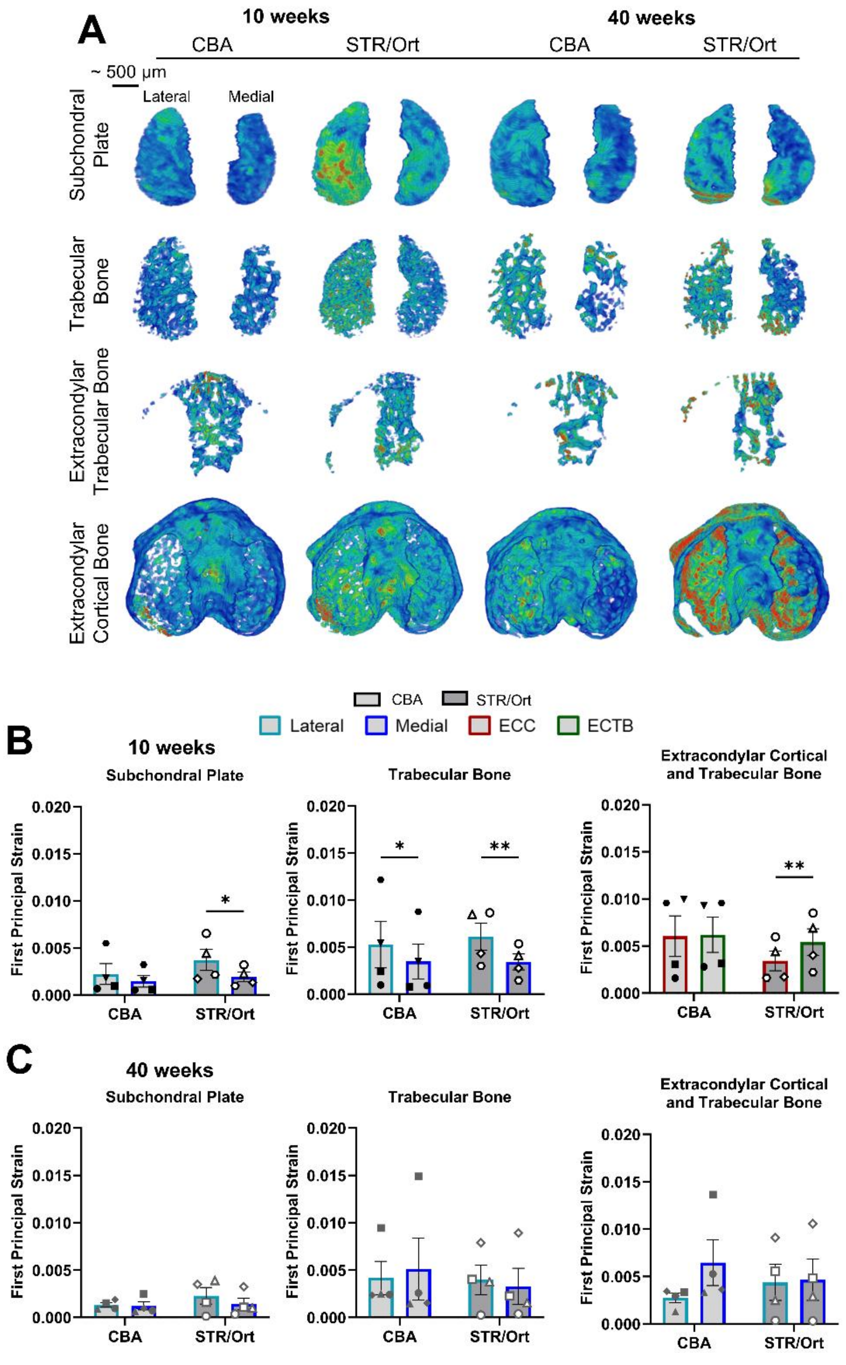


Supplementary Figure S4. Regionalised and condylar DVC-derived tensile strains are shown for the SCP, trabecular bone, extracondylar trabecular bone and extracondylar cortical bone from 10- and 40-week-old CBA and STR/Ort mice (A). Quantification of average tensile strain in the lateral (light blue outlined bar) and medial (dark blue outlined bar) condylar compartments of the tibial epiphyses of 10-week-old (B) and 40-week-old (C) mice show laterally-dominant accumulation of tension in 10-week-old STR/Orts, which is lost with ageing (D). In CBAs, tensile strains are evenly spread across lateral and medial condyles at both ages. Data are presented as mean ± SEM with symbols representing individual animals (N=4/age/genotype). Statistical significance between condyles was assessed using linear mixed model analyses with Fisher's LSD post-hoc test (C and D; *p<0.05, **p<0.01).

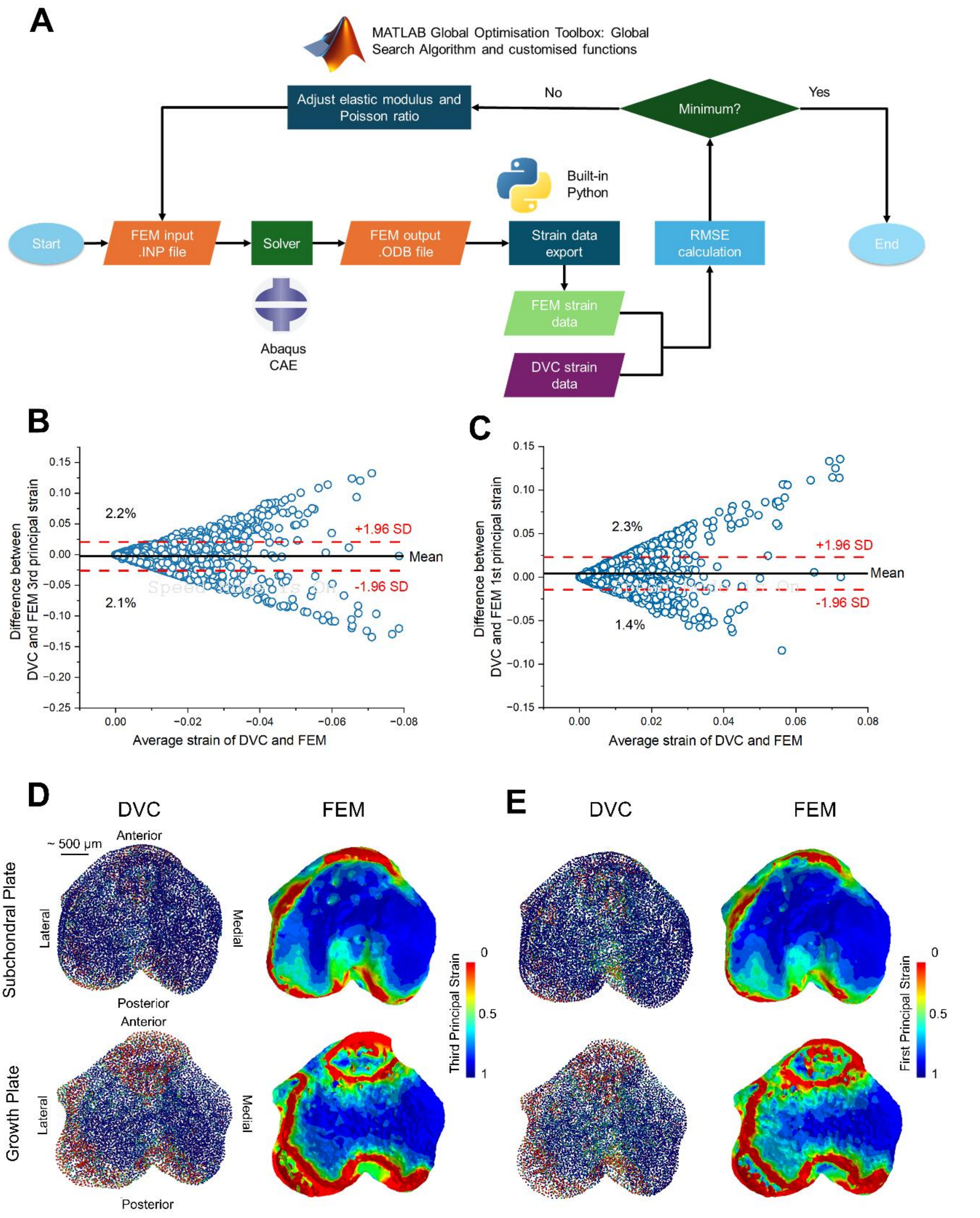


Supplementary Figure S5. Finite element model optimisation workflow and validation against DVC strain measurements. Schematic of iterative optimisation protocol used to refine finite element material properties (A). Elastic modulus and Poisson's ratio were systematically adjusted in MATLAB to minimise the RMSE between experimentally measured DVC strain fields and finite element predicted strain data. Bland-Altman plots comparing DVC and finite element derived compressive (B) and tensile (C) strains. Comparisons of DVC-derived compressive (D) and tensile (E) strains with DVC-optimised finite element-derived strains viewed from the SCP (top row) and growth plate (bottom row).

Supplementary Table S3. Quantification of regional thickness and volume in the tibial epiphyses of 10- and 40-week-old CBA and STR/Ort mice. Data presented as mean ± SEM for N=4 mice/age/genotype. Statistical significance was assessed using linear mixed model analysis with Fisher's LSD post-hoc test. † denotes statistical significance between 10- and 40-week-old animals, * denotes statistical significance between CBA and STR/Ort and § denotes statistical significance between anatomical region/condyle with single, double, triple and quadruple symbols corresponding to, $p<0.05$, $p<0.01$, $p<0.001$, and $p<0.0001$, respectively.

| | | | **Lateral Subchondral Plate** | **Medial Subchondral Plate** | **Lateral Trabecular Bone** | **Medial Trabecular Bone** | **Extracondylar Cortical Bone** | **Extracondylar Trabecular Bone** |
|---|---|---|---|---|---|---|---|---|
| **Thickness (µm)** | **10 weeks** | CBA | 73.06 ± 3.36 | 79.14 ± 1.56 § | 34.55 ± 0.71 | 34.72 ± 1.24 | 64.34 ± 1.97 | 28.8 ± 1.4 §§§§ |
| | | STR/Ort | 65.07 ± 3.73 | 83.52 ± 4.5 §§§ | 33. 42 ± 1.55 | 37.73 ± 1.03 §§ | 72.95 ± 1.69 ** | 31.01 ±1.82 §§§§ |
| | **40 weeks** | CBA | 82.03 ± 3.19 | 93.38 ± 5.79 † | 37.47 ± 0.92 | 36.35 ± 1.01 | 80.31 ± 1.94 †††† | 31.89 ± 0.85 §§§§ |
| | | STR/Ort | 79.01 ± 6.47 | 101.14 ± 2.47 §§, † | 37.95 ± 1.79 | 36.36 ± 3.12 | 85.11 ± 4.57 †† | 33.72 ± 0.83 §§§§ |
| **Volume ($mm^2$)** | **10 weeks** | CBA | 0.22 ± 0.006 | 0.16 ± 0.004 §§§§ | 0.18 ± 0.009 | 0.09 ± 0.002 §§§ | 0.95 ± 0.07 | 0.14 ± 0.01 §§§§ |
| | | STR/Ort | 0.19 ± 0.01 * | 0.2 ± 0.01 §§, ** | 0.2 ± 0.02 | 0.13 ± 0.007 §§, ** | 0.99 ± 0.07 | 0.15 ± 0.007 §§§§ |
| | **40 weeks** | CBA | 0.23 ± 0.04 | 0.28 ± 0.01 †† | 0.13 ± 0.01 †† | 0.07 ± 0.006 § | 1.28 ± 0.08 ††† | 0.09 ± 0.006 §§§§ |
| | | STR/Ort | 0.26 ± 0.01 †† | 0.21 ± 0.01 * | 0.13 ± 0.01 †† | 0.09 ± 0.02 † | 1.2 ± 0.07 †† | 0.09 ± 0.004 §§§§ |

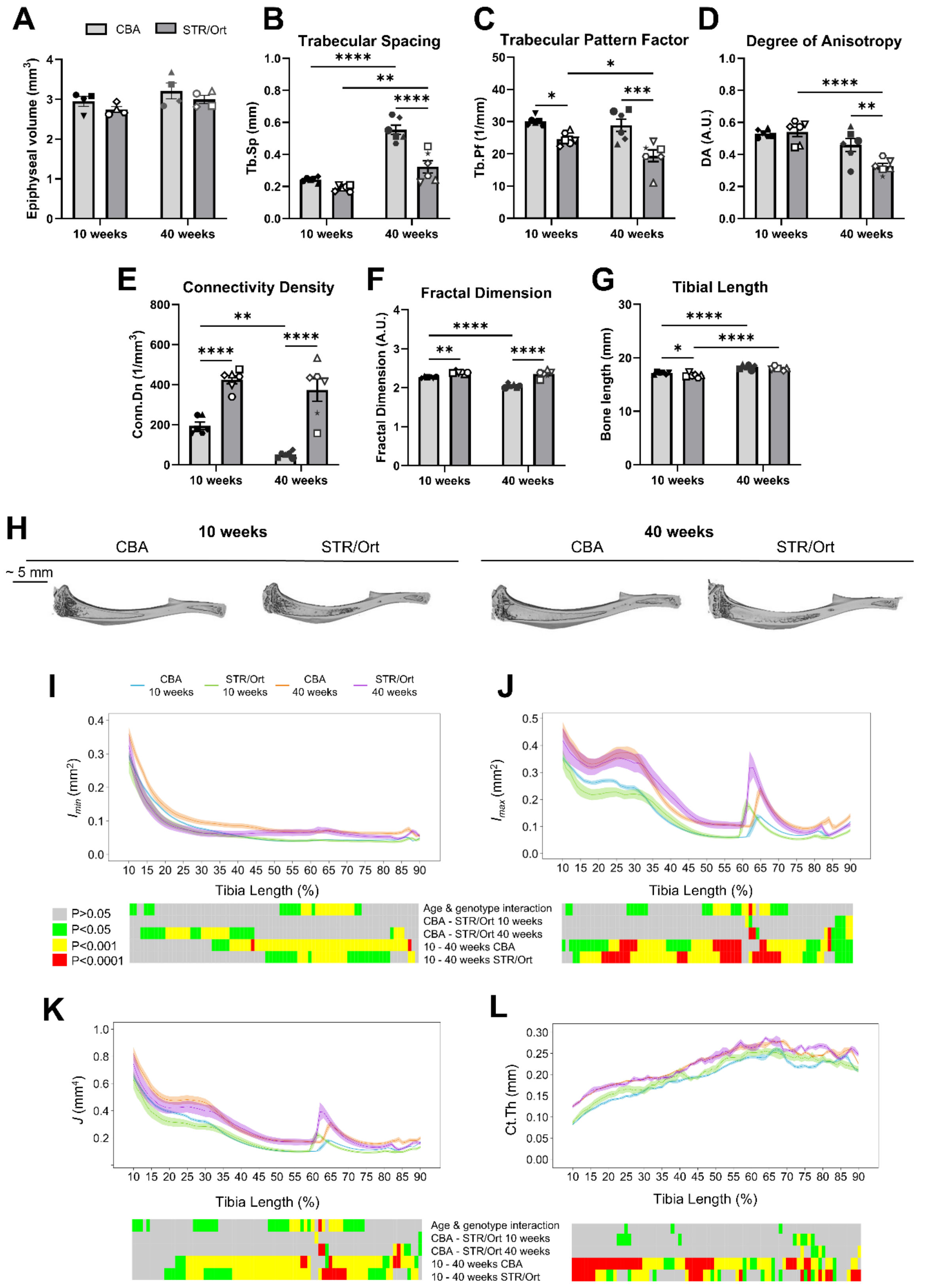


Supplementary Figure S6. MicroCT-based quantification of tibial epiphyseal volume (A) in addition to metaphyseal trabecular morphometric parameters including Tb.Sp (B), Tb.Pf (C), DA (D), Conn.D (E), fractal dimension (F) and tibial bone length (G) in 10- and 40-week-old CBA and STR/Ort mice. 3D rendering of tibial cortical bone from reconstructed microCT datasets show comparable anatomy between CBA and STR/Orts at both ages (H). Comparison of the $I_{min}$ (I), $I_{max}$ (J), J (K) and Ct.Th (L), evaluated between 10% and 90% of the tibial length are presented as line graphs representing mean ± SEM. Graphical heatmaps beneath show statistical differences at spatially matched locations along the tibial length between CBA and STR/Orts at 10- and 40-weeks of age. Data are presented as mean ± SEM with symbols representing individual animals (N=4/age/genotype, A or N=6/age/genotype, B-G). Statistical significance between groups was assessed using linear mixed model analyses with Fisher's LSD post-hoc test (A-H, *p<0.05, **p<0.01, ***p<0.001, and ****p<0.0001) and interaction between age and genotype was assessed by two-way ANOVA with Tukey's post-hoc test (J-M, grey, p>0.05; green, p<0.05; yellow, p<0.001; red, p<0.0001).
